# Self-healing codes: how stable neural populations can track continually reconfiguring neural representations

**DOI:** 10.1101/2021.03.08.433413

**Authors:** M. E. Rule, T. O’Leary

## Abstract

As an adaptive system, the brain must retain a faithful representation of the world while continuously integrating new information. Recent experiments have measured population activity in cortical and hippocampal circuits over many days, and found that patterns of neural activity associated with fixed behavioral variables and percepts change dramatically over time. Such “representational drift” raises the question of how malleable population codes can interact coherently with stable long-term representations that are found in other circuits, and with relatively rigid topographic mappings of peripheral sensory and motor signals. We explore how known plasticity mechanisms can allow single neurons to reliably read out an evolving population code without external error feedback. We find that interactions between Hebbian learning and single-cell homeostasis can exploit redundancy in a distributed population code to compensate for gradual changes in tuning. Recurrent feedback of partially stabilized readouts could allow a pool of readout cells to further correct inconsistencies introduced by representational drift. This shows how relatively simple, known mechanisms can stabilize neural tuning in the short term, and provides a plausible explanation for how plastic neural codes remain integrated with consolidated, long-term representations.

**Significance:** The brain is capable of adapting while maintaining stable long-term memories and learned skills. Recent experiments show that neural responses are highly plastic in some circuits, while other circuits maintain consistent responses over time, raising the question of how these circuits interact coherently. We show how simple, biologically motivated Hebbian and homeostatic mechanisms in single neurons can allow circuits with fixed responses to continuously track a plastic, changing representation without reference to an external learning signal.

---

The cellular and molecular components of the brain change continually. In addition to synaptic turnover (1), ongoing reconfiguration of the tuning properties of single neurons has been seen in parietal (2), frontal (3), visual (4, 5), and olfactory (6) cortices, and the hippocampus (7, 8). Remarkably, the “representational drift” (9) observed in these studies occurs without any obvious change in behavior or task performance. Reconciling dynamic reorganization of neural activity with stable circuit-level properties remains a major open challenge (9, 10). Furthermore, not all circuits in the brain show such prolific reconfiguration, including populations in primary sensory and motor cortices (11–13). How might populations with stable and drifting neural tuning communicate reliably? Put another way, how can an internally consistent ‘readout’ of neural representations survive changes in the tuning of individual cells?

These recent, widespread observations suggest that neural circuits can preserve learned associations at the population level while allowing the functional role of individual neurons to change (14–16). Such preservation is made possible by redundancy in population codes, because a distributed readout allows changes in the tuning of individual neurons to be offset by changes in others. However, this kind of stability is not automatic: changes in tuning must either be constrained in specific ways (e.g. 17, 18), or corrective plasticity needs to adapt the readout (19). Thus, while there are proposals for what might be required to maintain population codes dynamically, there are few suggestions as to how this might be implemented with known cellular mechanisms and without recourse to external reference signals that re-calibrate population activity to behavioral events and stimuli.

In this paper we show that the readout of continuous behavioral variables can be made resilient to ongoing drift as it occurs in a volatile encoding population. Such resilience can allow highly plastic circuits to interact reliably with more rigid representations. In principle, this permits compartmentalization of rapid learning to specialized circuits, such as the hippocampus, without entailing a loss of coherence with more stable representations elsewhere in the brain.

We provide a simple hierarchy of mechanisms that can tether stable and unstable representations using simple circuit architectures and well known plasticity mechanisms, Hebbian learning and homeostatic plasticity. Homeostasis is a feature of all biological systems, and examples of homeostatic plasticity in the nervous system are pervasive (e.g. 20, 21 for reviews). Broadly, homeostatic plasticity is a negative feedback process that maintains physiological properties such as average firing rates (e.g. 22, 23), neuronal variability (e.g. 24), distributions of synaptic strengths (e.g. 25, 26), and population-level statistics (e.g. 27, 28).

Hebbian plasticity complements homeostatic plasticity by strengthening connectivity between cells that undergo correlated firing, further reinforcing correlations (29, 30). Pairwise correlations in a population provide local bases for a so-called task manifold in which task-related neural activity resides (31). Moreover, neural representations of continuous variables typically exhibit bump-like single cell tuning that tiles variables redundlantly across a population (2, 7). We show how these features can be maintained to form a drifting population. We then show how Hebbian and homeostatic mechanisms can cooperate to allow a readout to track encoded variables despite drift, resulting in a readout that ‘self-heals’. Our findings thus emphasize a role for Hebbian plasticity in maintaining associations, as opposed to learning new ones.

Finally, we show how evolving representations can be tracked over substantially longer periods of time if a readout population encodes a stable predictive model of the variables being represented in a plastic, drifting population. Our assumptions thus take into account, and may reconcile, evidence that certain circuits and sub-populations maintain stable responses, while others, presumably those that learn continually, exhibit drift (32).

## Background

We briefly review representational drift and important recent work related to the ideas in this manuscript. Representational drift refers to ongoing changes in neural responses during a habitual task that are not associated with behavioural change (9). For example, in Driscoll et al. (2) mice navigated to one of two endpoints in a T-shaped maze (Fig. 1a), based on a visual cue. Population activity in Posterior Parietal Cortex (PPC) was recorded over several weeks using fluorescence calcium imaging. Neurons in PPC were tuned to the animal’s past, current, and planned behavior. Gradually, the tuning of individual cells changed: neurons could change the location in the maze in which they fired, or become disengaged from the task (Fig. 1b). The neural population code eventually reconfigured completely (Fig. 1c). However, neural tunings continued to tile the task, indicating stable task information at the population level. These features of drift have been observed throughout the brain (4, 5, 8). The cause for such ongoing change remains unknown. It may reflect continual learning and adaptation that is not directly related to the task being assayed, or unavoidable biological turnover in neural connections.

**Figure 1:**
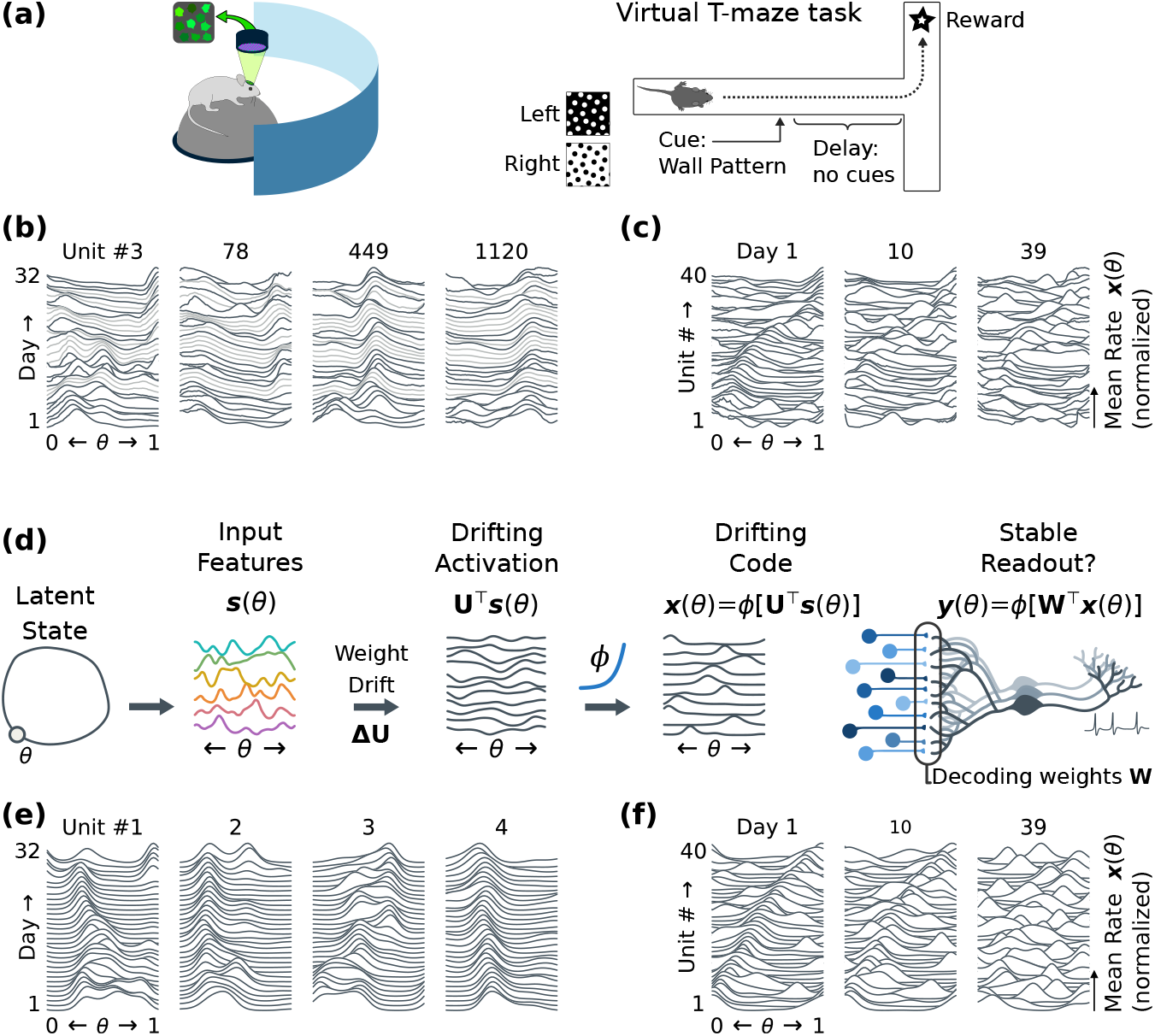
A model for representational drift. **(a)** Driscoll et al. (2) imaged population activity in PPC for several weeks, after mice had learned to navigate a virtual T-maze. Neuronal responses continued to change even without overt learning. **(b)** Tunings were often similar between days, but could change unexpectedly. Plots show average firing rates as a function of task pseudotime (0=beginning, 1=complete) for select cells from (2). Tuning curves from subsequent days are stacked vertically, from day 1 up to day 32. Missing days (light gray) are interpolated. Peaks indicate that a cell fired preferentially at a specific location (Methods: *Data and analysis*). **(c)** Neuronal tunings tiled the task. Within each day, the mouse’s behavior could be decoded from population activity (2, 19). Plots show normalized tuning curves for 40 random cells, stacked vertically. Cells are sorted by their preferred location on day 1. By day 10, many cells have changed tuning. Day 39 shows little trace of the original code. **(d)** We model drift in a simulated rate network (Methods: *Simulated drift*). An encoding population **x**(*θ*) receives input **s**(*θ*) with low-dimensional structure, in this case a circular track with location θ. The encoding weights **U** driving the activations **U**^⊤^**s**(θ) of this population drift, leading to drifting activations. Homeostasis preserves bump-like tuning curves. **(e)** As in the data (a-c), this model shows stable tuning punctuated by large changes. **(f)** The neural code reorganizes, while continuing to tile the task. We will examine strategies that a downstream readout could use to update how it decodes **x**(*θ*) to keep its own representation **y**(*θ*) stable. This readout is also modeled as linear-nonlinear rate neurons, with decoding weights **W**.

Previous work shows how downstream readouts could track gradual drift using external error feedback to re-learn how to interpret an evolving neural code, e.g. during ongoing rehearsal (19). Indeed, simulations confirm that learning in the presence of noise can lead to a steady state, in which drift is balanced by error feedback (33–36). Previous studies have shown that stable functional connectivity could be maintained despite synaptic turnover (33, 37, 38). More recent work has also found that discrete representations can be stabilized using neural assemblies that exhibit robust, all-or-nothing reactivation (39, 40).

Our work extends these results as follows. Rather than using external learning signals (19, 33–35), we show that drift can be tracked using internally generated signals. We allow the functional role of neurons in an encoding population to reconfigure completely, rather than just the synaptic connectivity (33, 37, 38). Previous work has explored how to stabilize point attractors using neuronal assemblies, both with stable (39, 41) and drifting (40) single-cell tunings. Key insights in this previous work include the idea that random reactivation or systematic replay of circuit states can reinforce existing point attractors. However, to plausibly account for stable readout of drifting sensorimotor information and other continuous variables observed experimentally (2, 7, 42), we require a mechanism that can track a continuous manifold, rather than point attractors.

Our other main contribution is to relate these somewhat abstract and general ideas to a concrete observations and relevant assumptions about the nature of drift. Representational drift is far from random (19) and this fact can be exploited to derive stable readouts. Specifically, the topology and coarse geometry of drifting sensorimotor representations appear to be consistent over time, while their embedding in population activity continually changes (2, 7, 43, 44). Thus the statistical structure of external variables is preserved, but not their neuron-wise encoding. Notably, brain-machine interface decoders routinely confront this, and apply online recalibration and transfer learning to track drift in (e.g. 45; 46 for review). We argue that neural circuits may do something similar to maintain ‘calibration’ between relatively stable circuits and highly plastic circuits. Early sensory or late motor populations that communicate directly with sensory receptors or muscle fibers necessarily have a consistent mapping between neural activity and signals in the external world. Such brain areas need to communicate with many other brain areas, including circuits that continually learn and adapt, and thus possess more malleable representations of behavioral variables.

## Results

We first construct a model of representational drift, in which homeostatic plasticity stabilizes the capacity of a “drifting” population to encode a continuous behavioural variable despite instability in single-neuron tunings. We then derive plasticity rules that allow single downstream neurons to stabilize their own readout of this behavioural variable despite drifting activity. Finally, we extend these ideas to show how comparatively stable neural populations that encode independent, predictive models of behavioural variables can actively track and stabilize a readout of drifting neural code.

### A model for representational drift

We have previously used the data from Driscoll et al. (47) to assess how much plasticity would be required to track drift in a linear readout (19). However, these data contain gaps of several days, and the number of high signal-to-noise units tracked for over a month is limited. To explore continual, long-term drift, we therefore construct a model inspired by the features of representational drift seen in spatial navigation tasks (2, 7).

We focus on key properties of drift seen in experiments. In both (2) and (7), neural populations encode continuous, lowdimensional behavioral variables (e.g. location), and exhibit localized ‘bump-like’ tuning to these variables. Tuning curves overlap, forming a redundant code. Over time, neurons change their preferred tunings. Nevertheless, on each day there is always a complete ‘tiling’ of a behavioral variable, thus the ability of the population to encode task information is conserved.

To model this kind of drifting code, we consider a population of *N* neurons that encode a behavioral variable, *θ*. We assume *θ* lies on a low-dimensional manifold, and is encoded in the vector of firing rates in a neural population with tuning curves **x**_*d*_(*θ*)={*x*_*d*,i_(*θ*),.,*x_d,N_*(*θ*)}^⊤^. These tunings change over time (day *d*).

We abstract away some details seen the experimental data in Fig. 1c. We focus on the slow component of drift, and model excess day-to-day tuning variability via a configurable parameter. We assume uniform coverage of the encoded space, which can be ensured by an appropriate choice of coordinates. We consider populations of 100 units that encode *θ*, and whose tunings evolve independently. Biologically, noise correlations and fluctuating task engagement would limit redundancy, but this would be offset by the larger number of units available.

To model drift, we first have to model an encoding ‘feature’ population whose responses depend on *θ*, and from which it is possible to construct bump-like tuning with a weighted readout (Fig. 1d). To keep our assumptions general, we do not assume that the encoding population has sparse, bump-like activity, and simply define a set of *K* random features (tuning curves), sampled independently from a random Gaussian process on *θ*. These features have an arbitrary but stable relationship to the external world, from which it is possible to reconstruct *θ* by choosing sufficiently large *K*:

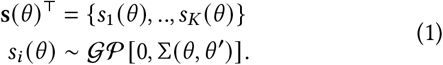

In the above equations, ∑(*θ*, *θ*′) denotes the covariance between the values of **s**(*θ*) at two states *θ* and *θ*′.

We next define an encoding of *θ* driven by these features with a drifting weight matrix **U**_*d*_={**u**_*d*,1_,., **u**_*d,N*_}, where 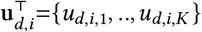 reflects the encoding weights for unit *x_d,i_*(*θ*) on day *d*. Each weight *u_d,i,j_* evolves as a discrete-time Ornstein-Uhlenbeck (OU) process, taking a new value on each day (Methods: *Simulated drift*). The firing rate of each encoding unit is given as a nonlinear function of the synaptic activation 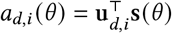:

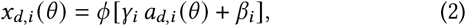

where *γ_i_* and *β_i_* are vectors that set the sensitivity and threshold of each unit. To model the nonlinear response of the readout and prevent negative firing rates, we use an exponential nonlinearity *ϕ*(·) = exp(·).

In this model, the mean firing-rate and population sparsity of the readout can be tuned by varying the sensitivity *γ* and threshold *β* in Eq. (2), respectively. In the brain, these single cell properties are regulated by homeostasis (24). Stabilizing mean rates 〈*x_d,i_*(*θ*)〉_*θ*_ ≈ *μ*_0_ ensures that neurons remain active. Stabilizing rate variability 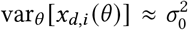 controls population code sparsity, ensuring that **x**_*d*_(*θ*) carries information about *θ* (48). This is achieved by adapting the bias *β_i_* and gain *γ_i_* of each unit *x_d,i_*(*θ*) based on the errors *ε_μ_*, *ε_σ_* between the statistics of neural activity and the homeostatic targets *μ*_0_, *σ*_0_:

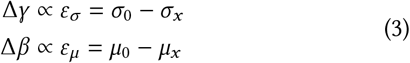

Fig. 1 shows that this model qualitatively matches the drift seen *in vivo* (2). Tuning is typically stable, with intermittent changes (Fig. 1e). This occurs because the homeostatic regulation in Eq. (3) adjusts neuronal sensitivity and threshold to achieve a localized, bump-like tuning curve at the location of peak synaptic activation, *θ*_0_. Changes in tuning arise when the drifting weight matrix causes the encoding neuron to be driven more strongly at a new value of *θ*. The simulated population code reconfigures gradually and completely over a period of time equivalent to several weeks in the experimental data (Fig. 1f).

### Hebbian homeostasis improves readout stability without external error feedback

Neural population codes are often redundant, with multiple units responding to similar task features. Distributed readouts of redundant codes can therefore be robust to small changes in the tuning of individual cells. We explored the consequences of using such a readout as an internal error signal to retrain synaptic weights in a readout population, thereby compensating for gradual changes in a representation without external feedback. This re-encodes a learned readout function **y**(*θ*) in terms of the new neural code **x**_*d*_(*θ*) on each “day” and improves the tuning stability of neurons that are driven by unstable population codes, even in single neurons. We first sketch an example of this plasticity, and then explore why this works.

Using our drifting population code as input, we model a readout population of *M* neurons with tuning curves **y**_*d*_(*θ*) = {*y*_*d*,1_(*θ*),.,*y_d,M_*(*θ*)}^⊤^ (Fig. 1d). We model this decoder as a linear-nonlinear function, using decoding weights **W** and biases (thresholds) **b** (leaving dependence on the day *d* implicit):

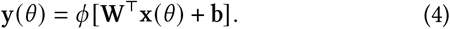

On each simulated “day”, we re-train the decoding weights using an unsupervised Hebbian learning rule (c.f. 49). This potentiates weights *w_i,j_* whose input *x_j_*(*θ*) correlates with the post-synaptic firing rate *y_i_*(*θ*). We modulate the learning rate by an estimate of the homeostatic error in firing-rate variability 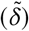. Thresholds are similarly adapted based on the homeostatic error in mean-rate 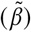. We include a small baseline amount of weight decay “*ρ*” and a larger amount of weight decay “*c*” that is modulated by 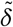. For a single readout neuron *y*(*θ*), the weights and biases evolve as:

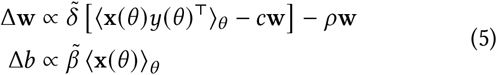

We apply Eq. (5) for 100 iterations on each simulated “day”, sampling all *θ* on each iteration. We assume that the timescale of Hebbian and homeostatic plasticity is no faster than the timescale of representational drift. The error terms 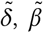 are leaky integrators of instantaneous errors (Eq. (3)) for each cell, *ε_σ_*, *ε_μ_*, respectively: 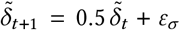 (analogously for 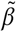, *ε_μ_*). For the readout **y**(*θ*), the homeostatic targets (*μ*_0_, *σ*_0_) are set to the firing-rate statistics in the initial, trained state (before drift has occurred). Eq. (5) therefore acts homeostatically. Rather than scale weights uniformly, it adjusts the component of the weights most correlated with the postsynaptic output, **y**(*θ*). Plasticity occurs only when homeostatic constraints are violated. Further discussion of this learning rule is given in Methods: *Synaptic learning rules*.

To test whether the readout can tolerate complete reconfiguration in the encoding population, we change encoding features one at a time. For each change, we select a new, random set of encoding weights **u***_i_* and apply homeostatic compensation to stabilize the mean and variability of *x_i_*(*θ*). Eq. (5) is then applied to update the decoding weights of the readout cell. This procedure is applied 200 times, corresponding to two complete reconfigurations of the encoding population of *N* =100 cells (Methods: *Single-neuron readout*).

With fixed weights, drift reduces the readout’s firing rate without changing its tuning (Fig. 2a),. This is because the initial tuning of the readout requires coincident activation of specific inputs to fire for its preferred *θ*_0_. Drift gradually destroys this correlated drive, and is unlikely to spontaneously create a similar conjunction of features for some other *θ*. For small amounts of drift, homeostasis Eq. (3) can stabilize the readout by compensating for the reduction in drive (Fig. 2b). Eventually, however, no trace of the original encoding remains. At this point, a new (random) *θ* will begin to drive the readout more strongly. Homeostasis adjusts the sensitivity of the readout to form a new, bump-like tuning curve at this location.

**Figure 2:**
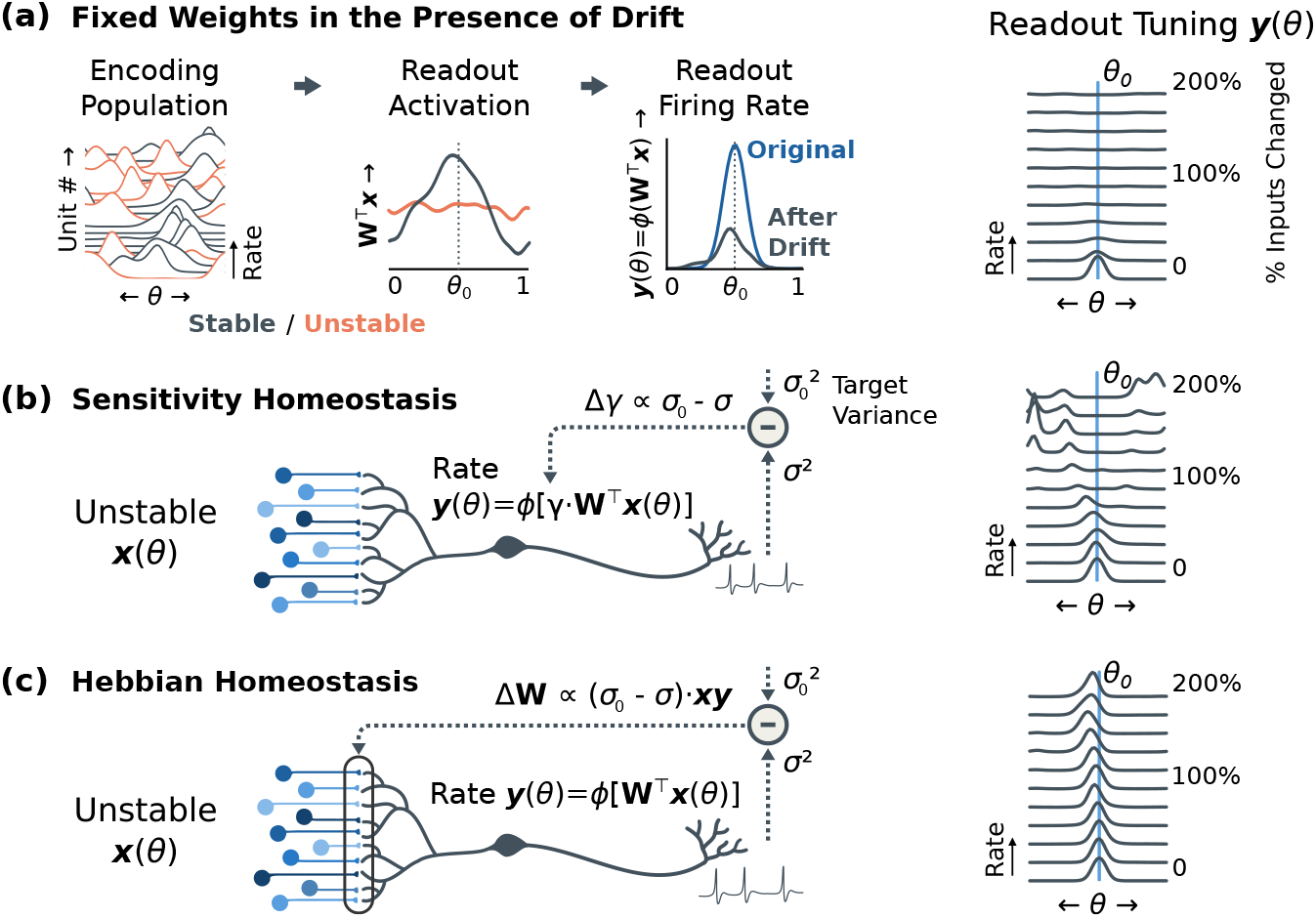
Homeostatic Hebbian plasticity enables stable readout from unstable populations. **(a)** Simulated drift in a redundant population causes a loss of excitability, but little change in tuning, to a downstream linear-nonlinear readout neuron. Since the cell is selective to a conjunction of features, it loses excitatory drive when some of its inputs change. Since most drift is orthogonal to this readout, however, the preferred tuning *θ*_0_ does not change. The right-most plot shows that the excitability diminishes as a larger fraction of inputs change. Two complete reconfigurations of an encoding population of 100 cells is shown. **(b)** Homeostatic adjustments to increase the readout’s sensitivity can compensate for small amounts of drift. As more inputs reconfigure, the cell compensates for loss of excitatory drive by increasing sensitivity (“gain”, *γ*). However, the readout changes to a new, random location once a substantial fraction of inputs have reconfigured (right). This phenomenon is the same as the model for tuning curve drift in the encoding population (c.f. Fig. 1e). **(c)** Hebbian homeostasis increases neuronal variability by potentiating synaptic inputs that are correlated with post-synaptic activity, or depressing those same synapses when neuronal variability is too high. This results in the neuron re-learning how to decode its own tuning curve from the shifting population code, supporting a stable readout despite complete reconfiguration (right). (Methods: *Single-neuron readout*)

Fig. 2c shows the consequences of Hebbian homeostasis (Eq. 5). Drift in the encoding **x**(*θ*) decreases the excitatory drive to the readout, activating Hebbian learning. Because small amounts of drift have minimal effect on tuning, the readout’s own output provides a self-supervised teaching signal. It relearns the decoding weights for inputs that have changed due to drift. Applying Hebbian homeostasis periodically improves stability, despite multiple complete reconfigurations of the encoding population. In effect, the readout’s initial tuning curve is transported to a new set of weights that estimate the same function from an entirely different input (for further discussion see Supplement: *Weight filtering*). In the long term the representation degrades, for reasons we dissect in the next section.

### Hebbian homeostasis with network interactions

In the remainder of the manuscript, we show how Hebbian homeostatic principles combine with population-level interactions to make readouts more robust to drift. Generally, a mechanism for tracking drift in a neural population should exhibit three features:

I. The readout should use redundancy to mitigate error caused by drift.
II. The readout should use its own activity as a training signal to update its decoding weights.
III. The correlations in input-driven activity in the readout neurons should be homeostatically preserved.

We explore three types of recurrent population dynamics that could support this: (1) Population firing-rate normalization; (2) Recurrent dynamics in the form of predictive feedback; (3) Recurrent dynamics in the form of a linear-nonlinear map. Fig. 5 summarizes the impact of each of these scenarios on a nonlinear population readout, and we discuss each in depth in the following subsections.

**Figure 3:**
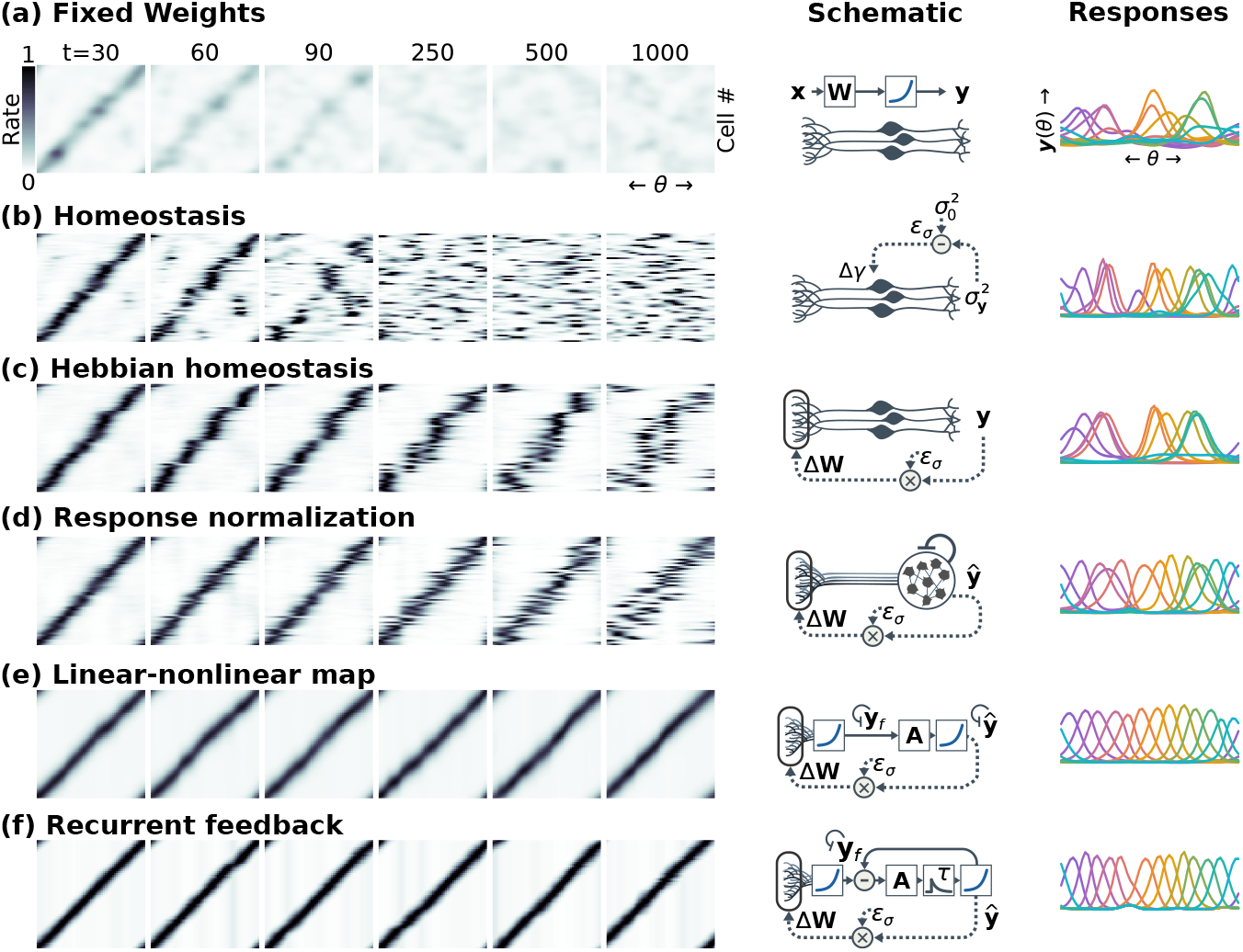
Self-healing in a nonlinear rate network. Each plot shows **(left)** a population readout **y**_0_(*θ*) from a drifting code **x**(*θ*) of *N* =100 cells; **(middle)** a schematic of the readout dynamics; and **(right)** a plot of readout tuning after applying each learning rule if 60% of the encoding cells were to change to a new tuning (Methods: *Population simulations*). **(a)** Drift degrades a readout with fixed weights. Drift is gradual, with *τ*=100. The simulated time frame corresponds to 10 complete reconfigurations. **(b)** Homeostasis increases sensitivity to compensate for loss of drive, but cannot stabilize tuning (*σ*^2^: firingrate variance, 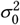: target variance, *ε_σ_*: homeostatic error, Δ*γ*: gain adjustment). **(c)** Hebbian homeostasis (Eq. 5) restores drive using the readout’s output as a training signal. Error correction is biased toward *θ* that drive more variability in the encoding population. (Δ**W**: weight updates) **(d)** Response normalization (Eq. 6) stabilizes the population statistics, but readout neurons can swap preferred tunings. (ŷ: normalized response). **(e)** A recurrent linear-nonlinear map (Eq. 8) pools information over the population, improving error correction (y_*f*_: feed-forward estimates, **A**: recurrent weights). **(f)** Predictive coding (Eq. 7) corrects errors via negative feedback (*τ* indicates dynamics in time). All simulations added 5% daily variability to **x**(*θ*), and applied 1% daily drift to the decoding weights **W**. Supplemental figure S2 evaluates readout stability for larger amounts ofvariability and readout-weight drift; Supplemental figure S3 quantifies readout stability for each scenario.

**Figure 4:**
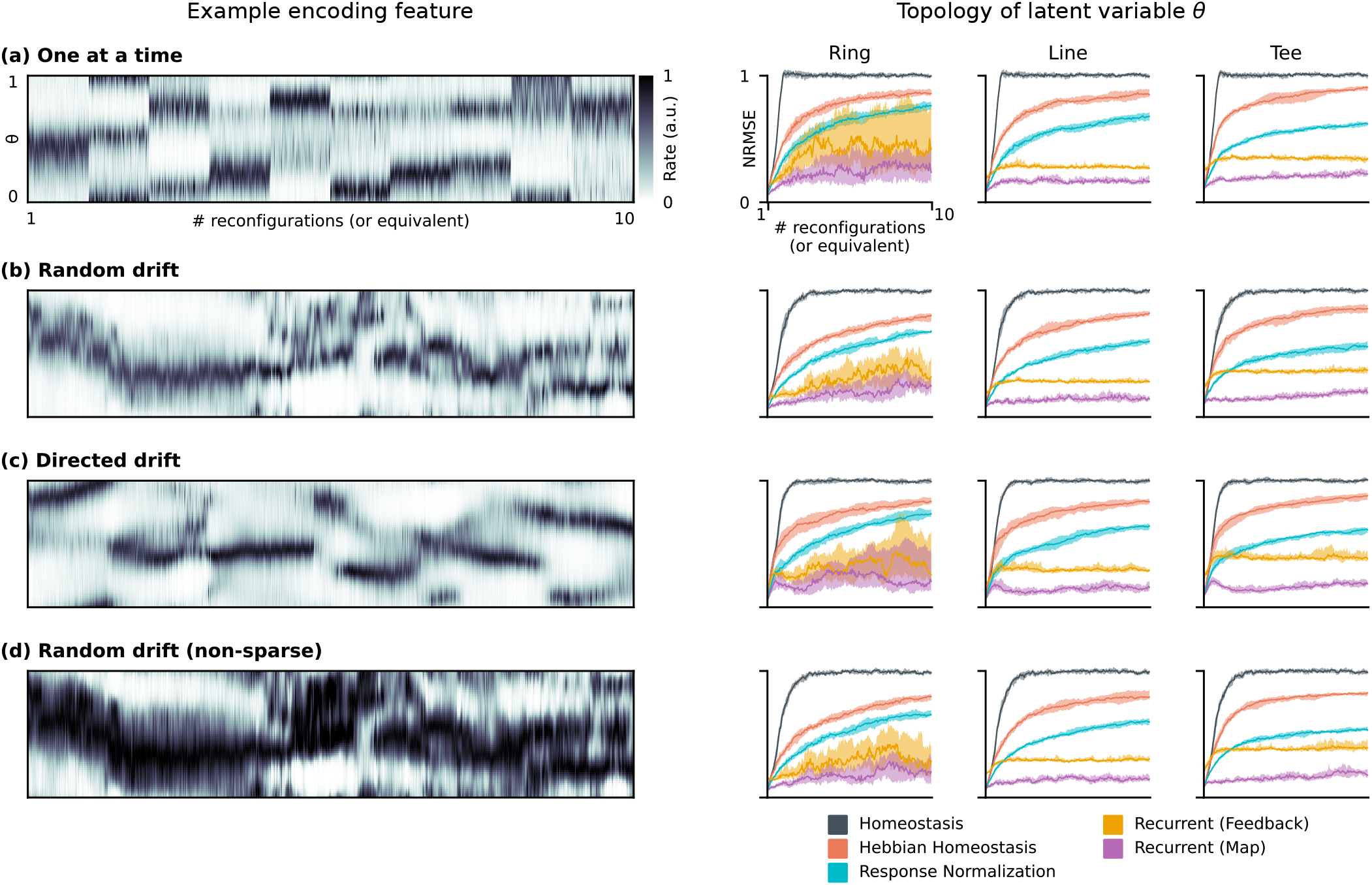
Self-healing stabilizes readout population codes for diverse types of representational drift. **Left:** An example of a single drifting encoding feature *x*(*θ*) sampled for a circular *θ* for different hypothetical drift scenarios. The horizontal axes for all plots are expressed in terms of the number of complete reconfigurations of the encoding population-code (or equivalent, for scenarios b-d). All simulations were run for *T*=1000 time-steps, corresponding to 10 complete reconfigurations in the encoding population. For continuous drift (b-d), time-constants were set to match the rate of population drift corresponding to changing encoding features one-at-a-time for a population of *N* = 100 encoding neurons. **Right:** Normalized Root-Mean-Squared-Error (NRMSE; 0=perfect match, 1=chance) of the readout population code over time. Lines indicate the median over 10 random seeds, and shaded regions the inter-quartile range. Simulation parameters are the same as for scenarios b-f in Figure 4 in the main text. We ran “self-healing” reconsolidation every Δ = 5 iterations (equivalent to 5% change in the encoding). We explored three topologies for *θ*: circular, linear, and T-maze (compare to Supplemental Figure S5). The rate of decay of the readout population code does not depend on the style of drift in the encoding population. **(a)** “One-at-a-time” drift changes one out of *N* = 100 encoding neurons on each iteration of the simulation. 100 simulated time-steps corresponds to one complete reconfiguration of the encoding population. **(b)** “Random drift” applies Ornstein-Uhlenbeck (OU) drift with a time constant *τ* = 100 (Methods: *Simulated drift*). **(c)** “Non-sparse drift” samples the encoding curves directly from a linear, Gaussian process, and does not apply the firing-rate nonlinearity *ϕ*(·). These features lack the sparse, bump-like tuning curves present in the other scenarios. The variance has been scaled to match that of the other drift scenarios. The correlation time is *τ* = 100. **(d)** Directed” drift simulates a second-order OU process evolving as two stages of Eq. (9) in the main text chained in series, such that consecutive changes are correlated in time. Each stage has a time constant *τ* = 50. Note that the circular environment is generally less stable than the liner and T-maze environments, since it is able to drift along a continuous degree of symmetry.

**Figure 5:**
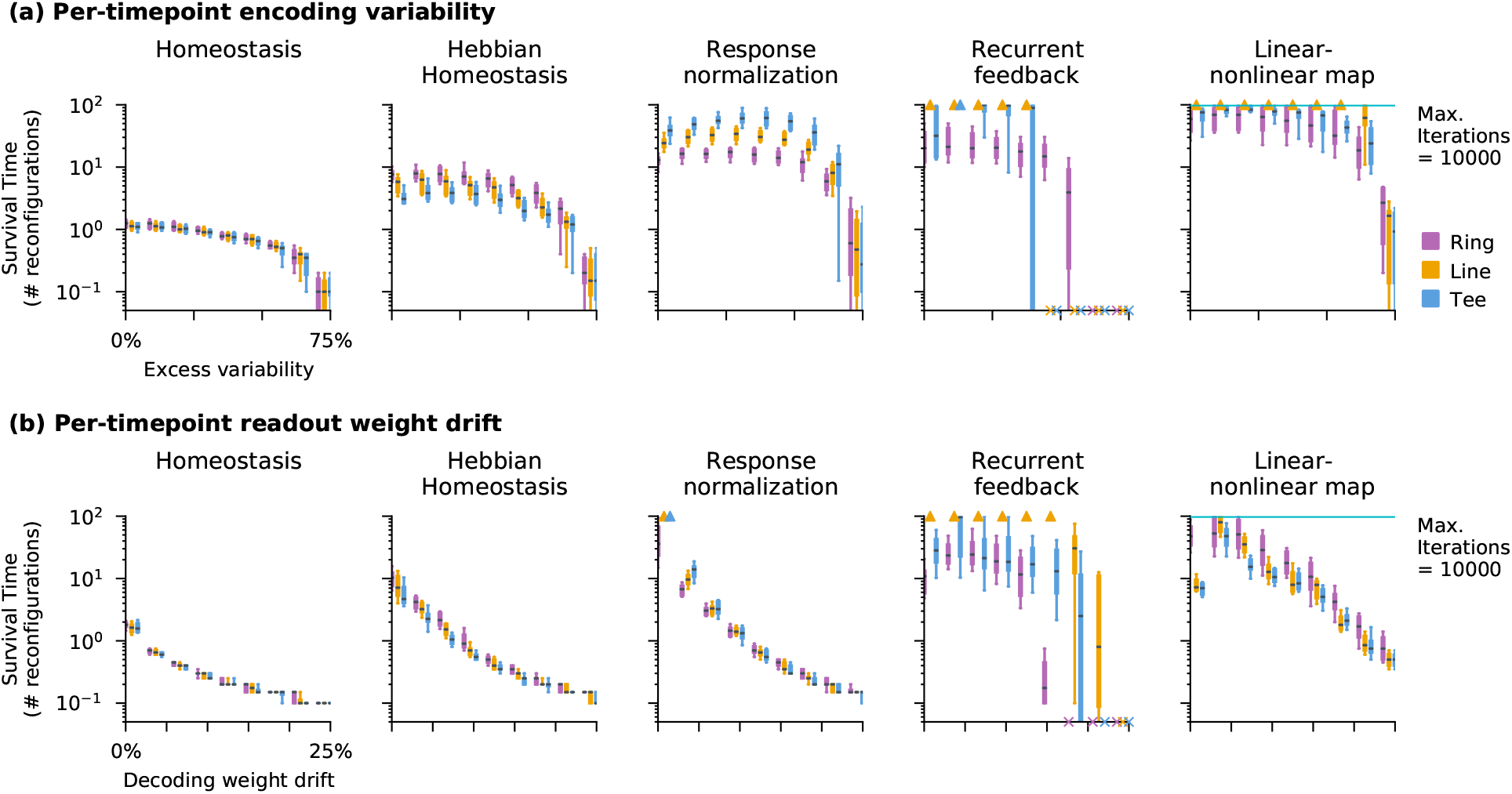
Encoding variability and readout-weight drift affect stability. Each plot shows the survival time for a self-healing readout as in Fig. 4 of the main text, measured as the time until the normalized root-mean-squared error of the readout exceeded 0.75. 10000 time-points are simulated for an encoding population of *N*=100 cells with a drift time constant of *τ*=100. Time (vertical axis) is expressed in multiples of the drift time constant. Boxes show the median (black) and inter-quartile range. Whiskers indicate the 10^th^-90^th^ percentiles over 20 random seeds. Scenarios in which the median survival time was less than one complete reconfiguration are plotted as “×” and those in which the median survival time exceeded the simulation time are are plotted as triangles. We explored three topologies for *θ*: circular, linear, and T-maze (compare to Supplemental Figure S5). All simulations used the same parameters as in Figure 4 in the main text, with the exception of the noise parameter (*r* or *u*) which is varied along the horizontal axis. Drift is gradual as described in Methods: *Simulated drift*. “Self-healing” reconsolidation is applied every Δ=5 time-steps. **(a)** We varied the amount of daily encoding variability that is unrelated to cumulative drift. This is expressed as the percentage of the variance in synaptic activation for the encoding neurons that is unique to each day (*r*, horizontal axis; Eq. (10) in the main text). The “response normalization” and “linear-nonlinear map” scenarios show good stability up until *u*≈40% (c.f. Fig. 4 d-f in the main text). “Recurrent feedback” is susceptible to instability, and some environments are destabilized at lower levels of variability. **(b)** We varied the amount of drift applied to the readout’s decoding weights **W** on each day (*n*, horizontal axis; Eq. (11) in the main text). Recurrent dynamics can correct small amounts of readout-weight drift, but stability degrades if drift exceeds ≈8% per time-step.

### Population competition with unsupervised Hebbian learning

In Fig. 2c, we saw that Hebbian homeostasis improved stability in the short term. Eq. (5) acts as an unsupervised learning rule, and pulls the readout *y*(*θ*) towards a family of bump-like tuning curves that tile *θ* (36). Under these dynamics, only drift Δ**x**(*θ*) that changes the peak of y(*θ*) to some new, nearby 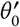 can persist. All other modes of drift are rejected. If the encoding population is much larger than the dimension of *θ*, there is large null space in which drift does not change the preferred tuning. However, in the long run Hebbian homeostasis drives the neural population toward a steady-state which forgets the initial tuning (Fig. 5c). This is because Hebbian learning is biased towards a few salient *θ*_0_ that capture directions in **x**(*θ*) with the greatest variability (30, 50, 51).

Models of unsupervised Hebbian learning address this by introducing competition among a population of readout neurons (50, 51). Such rules can track the full covariance structure of the encoding population, and lead to a readout population of bump-like tuning curves that tile the space *θ* (52–55). In line with this, we incorporate response normalization into a readout population (56). This serves as a fast-acting form of firing-rate homeostasis in Eq. (3), causing neurons to compete to remain active and encouraging diverse tunings (54, 57).

Because it is implemented via inhibitory circuit dynamics, we assume that this normalization acts quickly relative to plasticity, and model it by dividing the rates by the average firing rate across the population. If **y**_*f*_(*θ*) is the forward (unnormalized) readout from Eq. (4), we define the normalized readout **y**_*n*_(*θ*) by dividing out the average population rate, 〈**y**_*f*_(*θ*)〉_*M*_, and multiplying by a target mean rate *μ_p_*:

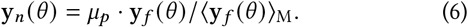

We found that response normalization improves readout stability (Fig. 5d). However, it does not constrain individual readout neurons to any specific preferred *θ*_0_. The readout thus remains sensitive to noise and perturbations, which in the long run can cause neurons to swap preferred tunings (Fig. 5d; Methods: *Population simulations*).

### Error-correcting recurrent dynamics

The error-correction mechanisms explored so far use redundancy and feedback to reduce errors caused by incremental drift. However, there is no communication between different readouts y_*i*_(*θ*) to ensure that the correlation structure of the readout population is preserved. In the remainder of the paper, we explore how a readout with a stable internal model for the correlation structure of **y**(*θ*) might maintain communication with a drifting population code.

Where might such a stable internal model exist in the brain? The dramatic representational drift observed in e.g. the hippocampus (7, 58) and parietal cortex (2) is not universal. Relatively stable tuning has been found in the striatum (59) and motor cortex (60–62). Indeed, perineuronal nets are believed limit structural plasticity in some mature networks (63), and stable connections can coexist with synaptic turnover (14). Drift in areas closer to the sensorimotor periphery is dominated by changes in excitability, which tends not to affect tuning preference of individual cells (3, 5). Thus, many circuits in the brain develop and maintain reliable population representations of sensory and motor variables. Moreover, such neural populations can compute prediction errors based on learned internal models (64), and experiments find that neural population activity recapitulates (65) and predicts (66) input statistics. Together, these findings suggest that many brain circuits can support relatively stable predictive models of the various latent variables that are used by the brain to represent the external world.

We therefore asked whether such a stable model of a behavioural variable could take the form of a predictive filter that tracks an unstable, drifting representation of that same variable. To incorporate this in our existing framework, we assume that the readout **y**(*θ*) contains a model of the “world” (*θ*) in its recurrent connections, which change much more slowly than **x**(*θ*). These recurrent connections generate internal prediction errors. We propose that these same error signals provide errorcorrection to improve the stability of neural population codes in the presence of drift.

We consider two kinds of recurrent dynamics. Both of these models are abstract, as we are not primarily concerned with the architecture of the predictive model, only its overall behavior. We first consider a network that uses inhibitory feedback to cancel the predictable aspects of its input, in line with models of predictive coding (67–69). We then consider a linear-nonlinear mapping that provides a prediction of **y**(*θ*) from a partially corrupted readout.

### Recurrent feedback of prediction errors

Some theories propose that neural populations retain a latent state that is used to predict future inputs (67–69). This prediction is compared to incoming information to generate a prediction error, which is fed back through recurrent interactions to update the latent state. This is depicted in the schematic in (Fig. 5f). Here, we assume that the network contains a latent state “**z**” and predicts the readout’s activity, **ŷ** = *ϕ*(**z**), with firing-rate nonlinearity *ϕ* as defined previously. Inputs provide a feed-forward estimate **y**_*f*_(*θ*), which is corrupted by drift. The prediction error is the difference between **y**_*f*_(*θ*) and **ŷ**. The dynamics of **z** are chosen as:

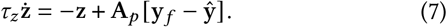

We set the weight matrix **A**_*p*_ to the covariance of the activations **z** = **W**^⊤^**x** during initial training (motivation for this choice is in Supplement: *Predictive coding as inference*). In making this choice, we assume that part of the circuit can learn and retain the covariance of **z**. This could in principle be achieved via Hebbian learning (49, 50, 70; Methods: *Learning recurrent weights*).

Assuming that a circuit can realise the dynamics in Eq. (7), the readout **ŷ** will be driven to match the forward predictions **y**_*f*_. We assume that this converges rapidly relative to the timescale at which **y**_*f*_(*θ*) varies. This improves the tracking of a drifting population code when combined with Hebbian homeostasis and response normalization (Fig. 5e). The readout continuously re-aligns its fixed internal model with the activity in the encoding population. We briefly discuss intuition behind why one should generally expect this to work.

The recurrent weights, **A**_*p*_, determine which directions in population-activity space receive stronger feedback. Feedback through larger eigenmodes of **A**_*p*_ is amplified, and these modes are rapidly driven to track **y**_*f*_. Due to the choice of **A**_*p*_ as the covariance of **z**, the dominant modes reflect directions in population activity that encode *θ*. Conversely, minor eigenmodes are weakly influenced by **y**_*f*_. This removes directions in population activity that are unrelated to *θ*, thereby correcting errors in the readout activity caused by drift.

In summary, Eq. (7) captures qualitative dynamics implied by theories of predictive coding. If neural populations update internal states based on prediction errors, then only errors related to tracking variations in *θ* should be corrected aggressively. This causes the readout to ignore “off manifold” activity in **ŷ**(*θ*) caused by drift. However, other models of recurrent dynamics also work, as we explore next.

### Low-dimensional manifold dynamics

Recurrent dynamics with a manifold of fixed-point solutions (distributed over *θ*) could also support error correction. We model this by training the readout to make a prediction **ŷ** of its own activity based on the feed-forward activity **y**_*f*_, via a linear-nonlinear map, (c.f. 71):

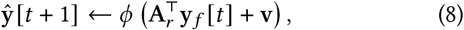

with timestep, *t*, and recurrent weights and biases **A**_*r*_ and **v**, respectively (Methods: *Learning recurrent weights*). We chose this discrete-time mapping for computational expediency, and Eq. (8) was applied once for each input *y_f_*(*θ*) alongside response normalization. In simulations, the recurrent mapping is also effective at correcting errors caused by drift, improving readout stability (Fig. 5f).

We briefly address some caveats that apply to both models of recurrent dynamics. The combination of recurrent dynamics and Hebbian learning is potentially destabilizing, because leaning can transfer biased predictions into the decoding weights. Empirically, we find that homeostasis (Eq. 3) prevents this, but must be strong enough to counteract all destabilizing influences. Additionally, when the underlying *θ* has continuous symmetries, drift can occur along these symmetries. This is evidenced by a gradual, diffusive rotation of the code for e.g. a circular environment. Other manifolds, like the T-shaped maze in (2), have no continuous symmetries and are not susceptible to this effect (Supplemental Figure S5). Overall, these simulations illustrate that internal models can constrain network activity. This provides ongoing error correction, preserves neuronal correlations, and allows neural populations to tolerate substantial reconfiguration of the inputs that drive them.

## Discussion

In this work, we derived principles that can allow stable and plastic representations to coexist in the brain. These self-healing codes have a hierarchy of components, each of which facilitates a stable readout of a plastic representation: (I) Singlecell tuning properties (bump-like tuning, redundant tiling of encoded variables) that make population codes robust to small amounts of drift; (II) Populations that use their own output as a training signal to update decoding weights, and (III) Circuit interactions that track evolving population statistics using stable internal models. All of these components are biologically plausible, some corresponding to single-cell plasticity mechanisms (Hebbian and homeostatic plasticity), others corresponding to circuit architectures (response normalization, recurrence) and others corresponding to higher-level functions that whole circuits appear to implement (internal models). As such, these components may exist to a greater or lesser degree in different circuits.

Hebbian plasticity is synonymous with learning novel associations in much of contemporary neuroscience. Our findings argue for a complementary hypothesis that Hebbian mechanisms can also reinforce learned associations in the face of ongoing change, in other words, prevent spurious learning. This view is compatible with the observation that Hebbian plasticity is a positive feedback process, where existing correlations become strengthened, in turn promoting correlated activity (72). Abstractly, positive feedback is required for hysteresis, which is a key ingredient of any memory retention mechanism, biological or otherwise, because it rejects external disturbances by reinforcing internal states.

Homeostasis, by contrast, is typically seen as antidote to possible runaway Hebbian plasticity (72). However, this idea is problematic due to the relatively slow timescale at which homeostasis acts (73). Our findings posit a richer role for homeostatic (negative) feedback in maintaining and distributing responsiveness in a population. This is achieved by regulating the mean and the variance of neural activity (24).

We considered two populations: a drifting population that encodes a behavioral variable, and another that extracts a drift-resilient readout. This could reflect communication between stable and plastic components of the brain, or the interaction between stable and plastic neurons within the same circuit. This is consistent with experiments that find consolidated stable representations (12, 16), or with the view that neural populations contain a mixture of stable and unstable cells (74).

By itself, Hebbian homeostasis preserves population codes in the face of drift over a much longer timescale than the lifetime of a code with fixed readout (Fig. 2). Even though this mechanism ultimately corrupts a learned tuning, the time horizon over which the code is preserved may be adequate in a biological setting, particularly in situations where there are intermittent opportunities to reinforce associations behaviourally. However, in the absence of external feedback, extending the lifetime of this code still further required us to make additional assumptions about circuit structures that remain to be tested experimentally.

We found that a readout population can use an internal model to maintain a consistent interpretation of an unstable encoding population. Such internal models are widely hypothesized to exist in various guises (64, 66, 67, 69). We did not address how these internal models are learned initially, nor how they might be updated. By setting fixed recurrent weights, we are also assuming that population responses in some circuits are not subject to drift. This may be reasonable, given that functional connectivity and population tuning in some circuits and subpopulations is found to be stable (11–13).

The recurrent architectures we studied here are reminiscent of mechanisms that attenuate forgetting via replay (e.g. 75, 76). The internal models must be occasionally re-activated through rehearsal or replay to detect and correct inconsistencies caused by drift. If this process occurs infrequently, drift becomes large, and the error correction will fail.

The brain supports both stable and volatile representations, typically associated with memory retention and learning, respectively. Artificial neural networks have so far failed to imitate this, and suffer from catastrophic forgetting wherein new learning erases previously learned representation (77). Broadly, most proposed strategies mitigate this by segregating stable and unstable representations into distinct subspaces of the possible synaptic weight changes (c.f. 18). These learning rules therefore prevent disruptive drift in the first place. The mechanisms explored here do not restrict changes in weights or activity: the encoding population is free to reconfigure its encoding arbitrarily. However, any change in the code leads to a complementary change in how that code is read out. Further exploration of these principles may clarify how the brain can be simultaneously plastic and stable, and provide clues to how to build artificial networks that share these properties.

## Materials and Methods

### Data and analysis

Data shown in Fig. 1b,c were taken from Driscoll et al. (2), and are available online at at Dryad (47). Examples of tuning curve drift were taken from mouse four, which tracked a sub-population of cells for over a month using calcium fluorescence imaging. Normalized logfluorescence signals (ln[*x*/〈*x*〉]) were filtered between 0.3 and 3 Hz (4^th^ Butterworth, forward-backward filtering), and individual trial runs through the T maze were extracted. We aligned traces from select cells based on task pseudotime (0: start, 1: reward). On each day, we averaged log-fluorescence over all trials and exponentiated to generate the average tuning curves shown in Fig. 1b. For Fig. 1c, a random subpopulation of forty cells was sorted based on their peak firing location on the first day. For further details, see (2, 19).

### Simulated drift

We modeled drift as a discrete-time Ornstein-Uhlenbeck (OU) random walk on encoding weights **U**, with time constant *τ* (in days) and perday noise variance *α*. We set the noise variance to *α*=2/*τ* to achieve unit steady-state variance. Encoding weights for each day are sampled as:

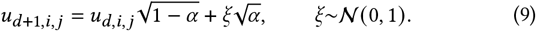

These drifting weights propagate the information about *θ* available in the features **s**(*θ*) (Eq. 1) to the encoding units **x**(*θ*), in a way that changes randomly over time.

This random walk in encoding-weight space preserves the population code statistics on average: It preserves the geometry of *θ* in the correlations of **a**_*t*_(*θ*), and the average amount of information about *θ* encoded in the population activations (Supplement: *Stability of encoded information*). This implies that the difficulty of reading out a given tuning curve *y*(*θ*) (in terms of the L2 norm of the decoding weights, ||**w**_*j*_||^2^) should remain roughly constant over time. This assumption, that **x**(*θ*) encodes a stable representation for θ in an unstable way, underlies much of the robustness we observe. We discuss this further in Methods: *Synaptic learning rules*.

Because the marginal distribution of the encoding weights on each day is Gaussian, 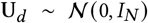, the synaptic activations 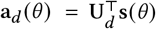 are samples from a Gaussian process on *θ*, with covariance inherited from **s**(*θ*) (Supplement: *Gaussian-process tuning curves*). In numerical experiments, we sampled the synaptic activation functions **a**_*d*_(*θ*) from this Gaussian process directly. We simulated *θ* ∈ [0, 1) over a discrete grid with 60 bins, sampling synaptic activations from a zero-mean Gaussian process on *θ* with a spatially-low-pass squared-exponential kernel (*σ* = 0.1). The gain and threshold (Eq. 2) for each encoding unit was homeostatically adjusted for a target meanrate of *μ*_0_ = 5 and rate variance of 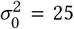 (in arbitrary units). This was achieved by running Eq. (3) for 50 iterations with rates *η_γ_*, = 0.1, *η_β_* = 0.2 for the gain and bias homeostasis, respectively.

To show that the readout can track drift despite complete reconfiguration of the neural code, we replace gradual drift in all features with abrupt changes in single features in Fig. 2. For this, we re-sampled the weights for single encoding units one-at-a-time from a standard normal distribution. Self-healing plasticity rules were run each time 5 out of the 100 encoding features changed. Supplemental Fig. S1 confirms that abrupt drift in a few units is equivalent to gradual drift in all units. Unless otherwise stated, all other results are based on an OU model of encoding drift.

We modeled excess variability in the encoding population that was unrelated to cumulative drift. This scenario resembles the drift observed in vivo (9; Supplemental Fig. S4). We sampled a unique “perday” synaptic activation *ã_d,i_*(*θ*) for each of the encoding units, from the same Gaussian process on *θ* used to generate the drifting activation functions *a_d,i_*(*θ*). We mixed these two functions with a parameter *r* = 0.05 such that the encoding variability was preserved (i.e. 5% of the variance in synaptic activation is related to random variability):

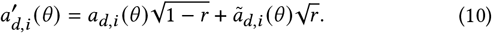

Supplemental Fig. S2a shows that the readout can tolerate up to 30% excess variability with modest loss of stability. Supplemental Fig. S4 shows that neuronal recordings from Driscoll et al. (47) are consistent with 30% excess variability, and that the qualitative conclusions of this paper hold for this larger amount of day-to-day variability (Supplement: *Calibrating the model to data*).

We also applied drift on the decoding synapses **W**. This is modeled similarly to Eq. (10), with the parameter *n* controlling the percentage of variance in synapse weight that changes randomly at the start of each each day:

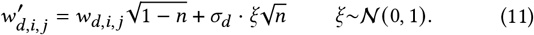

where *σ_d_* is the empirical standard-deviation of the the decoding weights on day *d*. Unless otherwise stated, we use *n* = 1%. Larger values of drift on the decoding weights is destabilizing for Hebbian homeostasis (with or without response normalization), but readouts with stable internal recurrent dynamics can tolerate larger (~ 8%) amounts of readout-weight drift (Supplemental Fig. S2b).

### Synaptic learning rules

The learning rule in Eq. (5) is classical unsupervised Hebbian learning, which is broadly believed to be biologically plausible (49, 50, 70). However, it has one idiosyncrasy that should be justified: The rates of learning and weight decay are modulated by a homeostatic error in firing-rate variability. The simplest interpretation of Eq. (5) is a premise or ansatz: learning rates should be modulated by homeostatic errors. This is a prediction that will need to be experimentally confirmed. Such a learning might be generically useful, since it pauses learning when firing-rate statistics achieve a useful dynamic range for encoding information. The fact that weight decay is proportional to learning rate is also biologically plausible, since each cells has finite resources to maintain synapses.

*Eq*. (5) may also emerge naturally from the interaction between homeostasis and learning rules in certain scenarios. When Hebbian learning is interpreted as a supervised learning rule, it is assumed that other inputs bias the spiking activity *y* of a neuron toward a target *y*^*^. This alters the correlations between presynaptic inputs **x** and post-synaptic spiking. Hebbian learning rules, especially temporally asymmetric ones based on spike timing (78), adjust readout weights **w** to potentiate inputs that correlate with this target. In the absence of external learning signals, homeostatic regulation implies a surrogate training signal *ỹ*^*^. This *ỹ*^*^ is biased toward a target mean-rate and selectivity. For example, recurrent inhibition could regulate both population firing rate and population-code sparsity. This could restrict postsynaptic spiking, causing Hebbian learning to adjust readout weights to achieve the desired statistics. Cells may also adjust their sensitivity and threshold homeostatically. Hebbian learning could then act to adjust incoming synaptic weights to achieve the target firing-rate statistics, but in a way that is more strongly correlated with synaptic inputs.

In Supplement: *Hebbian homeostasis as an emergent property*, we verify the intuition that Eq. (5) should arise through emergent interactions between homeostasis and Hebbian learning in a simplified, linear model. In the remainder of this section, we use a linear readout to illustrate why one should expect Eq. (5) to be stabilizing.

The decoder’s job is to generate a stable readout from the drifting code **x**(*θ*). This is a regression problem: the decoding weights **W** should map **x**(*θ*) to a target **y**(*θ*). Since small amounts of drift are akin to noise, **W** should be regularized to improve robustness. The L2-regularized linear least-squares solution for **W** is:

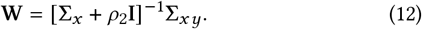

The regularization *ρ*_2_**I** corresponds to the assumption that drift will corrupt the activity of **x**(*θ*) by an amount 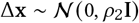.

Can drift be tracked by re-inferring **W** on each day? We lack the ground-truth covariance 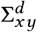 to re-train **W**, but could estimate it from decoded activity **y**_*d*_(*θ*):

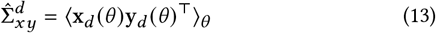

Since **y**_*d*_(*θ*) is decoded from **x**_*d*_(*θ*) through weights **W**_*d*_, the estimated covariance is 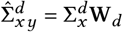, where 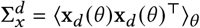 is the covariance of the inputs **x**_*d*_(*θ*). The regularized least-squares weight update is therefore:

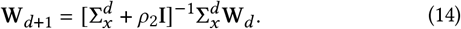

This update can be interpreted as recursive Bayesian filtering of the weights (Supplement: *Weight filtering*).

Because **x**(*θ*) continues to encode information about *θ*, we know that variability in the decoded **y**(*θ*) should be conserved. Each readout *y_i_*(*θ*) homeostatically adjusts its sensitivity to maintain a target variability 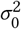. As introduced earlier, this multiplies the firing rate by a factor *γ* = *σ*_0_/*σ*_1_ = 1 **+** *δ*, where *δ* is a small parameter. and *σ*_1_ is the standard deviation of the readout’s firing rate after drift but before normalization. Accounting for this in Eq. (14) and considering the weight vector **w** for a single readout neuron yields:

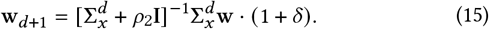

To translate this into a plausible learning rule, the solution Eq. (15) can be obtained via gradient descent. Recall the loss function 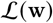 for optimizing regularized linear least-squares:

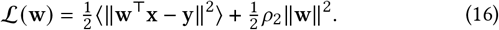

Gradient descent 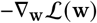 on Eq. (16) implies the weight update

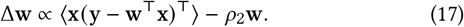

After matching terms between Eqs. (12–15) and Eq. (17) and simplifying, one finds the following Hebbian learning rule:

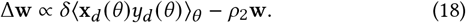

Eq. (18) is equivalent to Eq. (5) for a certain regularization strength *ρ*_2_ (now taking the form of weight decay). The optimal value of *ρ*_2_ depends on the rate of drift. Since drift drives homeostatic errors, it follows that *ρ*_2_ ∝ *δ* for small *δ*. Here, we set *ρ*_2_ = *δ*, corresponding to *c* = 1 in Eq. (18).

### Single-neuron readout

In Fig. 2, we simulated a population of 100 encoding neurons **x**_*d*_(*θ*) that changed one at a time (Methods: *Simulated drift*). We initialized a single readout *y*(*θ*) = *ϕ* [**w**^⊤^**x**(*θ*)] to decode a Gaussian bump *y*_0_(*θ*) (*σ* = 5% of the track length) from the activations **x**_0_(*θ*) on the first day. We optimized this via gradient descent using a linear-nonlinear Poisson loss function.

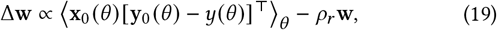

with regularizing weight decay *ρ_r_* = 10^-4^. In this deterministic firingrate model, the Poisson error allows the squared-norm of the residuals to be proportional to the rate. We simulated 200 time points of drift, corresponding to two complete reconfigurations of the encoding population. After each encoding-unit change, we applied 100 iterations of either naïve homeostasis (Fig. 2b; Eq. 3) or Hebbian homeostasis (Fig. 2c; Eq. 5). For naïve homeostasis, the rates for gain and threshold homeostasis were *η_β_* = 10^-3^ and *η_γ_* = 10^-5^, respectively. For Hebbian homeostasis, the rates were *η_β_* = 10^-1^ and *η_γ_* = 10^-3^.

Homeostatic regulation requires averaging the statistics over time (48). To model this, we calculated the parameter updates for the gain and bias after replaying all *θ* and computing the mean and variance of the activity for each neuron. Since the processes underlying cumulative changes in synaptic strength are also slower than the timescale of neural activity, weight updates were averaged over all *θ* on each iteration. We applied additional weight decay with a rate *ρ* = 1 × 10^-4^ for regularization and stability, and set *c* =1 in Eq. (5) such that the rate of weight decay was also modulated by the online variability error 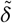.

### Learning recurrent weights

For recurrent dynamics modeled as feedback in Eq. (7), supervised, linear Hebbian learning implies that the recurrent weights should be proportional to the covariance of the state variables **z**. To see this, consider a linear Hebbian learning rule, where **z** has been entrained by an external signal, and serves as both the presynaptic input and postsynaptic output:

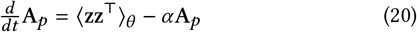

Where *α* is a weight decay term. This has a fixed point at **A**_*p*_ = 〈**zz**^⊤^)/*α*. In our simulations, we ensure that **z** is zero-mean such that the second moment, 〈**zz**^⊤^〉, is equal to the covariance.

For the linear-nonlinear map model of recurrent dynamics Eq. (8), neurons could learn **A**_*r*_ by comparing a target **y**_0_ to the predicted **y**_*r*_ at the same time that the initial decoding weights **W**_0_ are learned. For example, **y**_0_ could be an external (supervised) signal or the forward predictions in Eq. (4) before drift occurs, and **y**_*r*_ could arise though recurrent activity in response to **y**_0_. A temporally-asymmetric plasticity rule could correlate the error between these signals with the recurrent synaptic inputs to learn **A**_*r*_ (78). This plasticity rule should update weights in proportion to the correlations between synaptic input 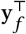 and a prediction error **y**_0_ – **y**_*r*_:

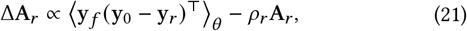

where *ρ_r_* = 10^-4^ sets the amount of regularizing weight decay.

Eq. (8) is abstract, but captures the two core features of error correction through recurrent dynamics. It describes a population of readout neurons that predict each-other’s activity through recurrent weights. Eq. (21) states that these weights are adapted during initial learning to minimize the error in this prediction. We assume **A**_*r*_ is fixed once learned.

### Population simulations

In Fig. 5, we simulated an encoding population of 100 units. Drift was simulated as described in Methods: *Simulated drift*, with *τ* = 100. In all scenarios, we simulated *M* = 60 readout cells tiling a circular *θ* divided into *L* = 60 discrete bins. Learning and/or homeostasis was applied every 5 iterations of simulated drift. The readout weights and tuning curves were initialized similarly to the single-neuron case, but with tuning curves tiling *θ*.

For the predictive coding simulations (Eq. 7), we simulated a second inner loop to allow the network activity **z** to reach a steady state for each input **x**(*θ*). This loop ran for 100 iterations, with time constant of *τ_z_* = 100. The recurrent weights **A**_*p*_ were initialized as the covariance of the synaptic activations on the first day (**Σ**_*z*_ where **z**(*θ*) = **W**^⊤^**x**(*θ*)) and held fixed over time. The final value **z** was used to generate a training signal, 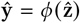, to update the readout weights. For the recurrent map, recurrent weights were learned initially using Eq. (21) and held fixed through the simulations.

For both the linear-nonlinear map and the recurrent feedback models, weights were updated as in Eq. (5), where the output of the recurrent dynamics was used to compute homeostatic errors and as the signal **ŷ** in Hebbian learning. For naïve homeostasis (Fig. 5b) and Hebbian homeostasis (with and without response normalization; Fig. 5cd), learning rates were the same as in the single-neuron simulations (Fig. 2; Methods; *Single-neuron readout*). For the linear-nonlinear map (Fig 5e), learning rates were set to *η_γ_* = 10^-4^ and *η_β_* = 10^-1^. For recurrent feedback (Fig 5f), the learning rates were *η_γ_* = 5 × 10^-3^ and *η_β_* = 5. Learning rates for all scenarios were optimized via grid search.

Response normalization was added on top of Hebbian homeostasis for Fig. 5d, and was also included in Fig. 5ef to ensure stability. The population rate target *μ_p_* for response normalization was set to the average population activity in the initially trained state.

Different parameters were used to generate the right-hand column of Fig. 5, to show the effect of a larger amount of drift. After training the initial readout, 60% of the encoding features were changed to a new, random tuning. Learning rates were increased by 50× for naïve homeostasis to handle the larger transient adaptation needed for this larger change. The other methods did not require any adjustments in parameters. Each homeostatic or plasticity rule was then run to steady-state (1000 iterations).

**Figure 6:**
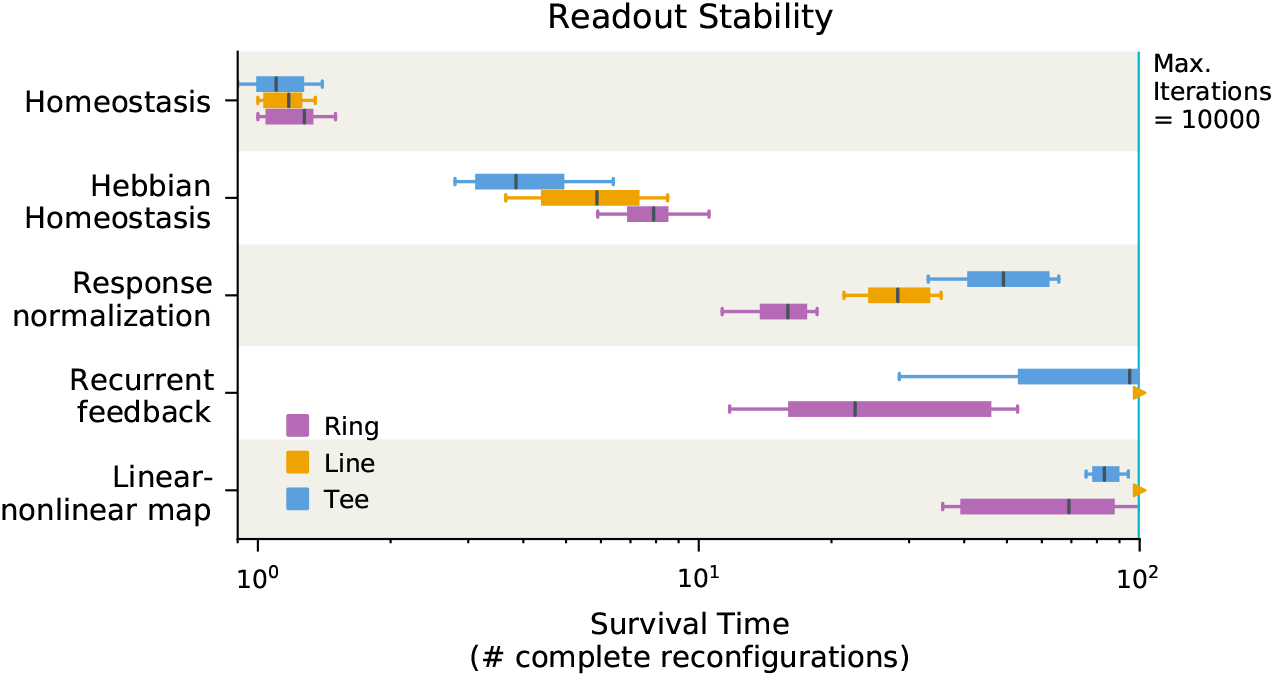
Summary of stability of each model scenario. Each box shows survival time for a self-healing readout as in Fig. 4 of the main text, measured as the time until the normalized root-mean-squared error of the readout exceeded 0.75. 10000 time-points are simulated for an encoding population of *N*=100 cells with a drift time constant of *τ* =100. Time (horizontal axis) is expressed in multiples of the drift time constant. Boxes show the median (black) and inter-quartile range. Whiskers indicate the 10^th^-90^th^ percentiles over 20 random seeds. Scenarios in which the median survival time exceeded the simulation time are are plotted as triangles. We explored three topologies for *θ*: circular, linear, and T-maze (compare to Supplemental Figure S5). All simulations used the same parameters as in Figure 4 in the main text. Drift is gradual as described in Methods: *Simulated drift*. “Self-healing” reconsolidation is applied every Δ=5 time-steps.

**Figure 7:**
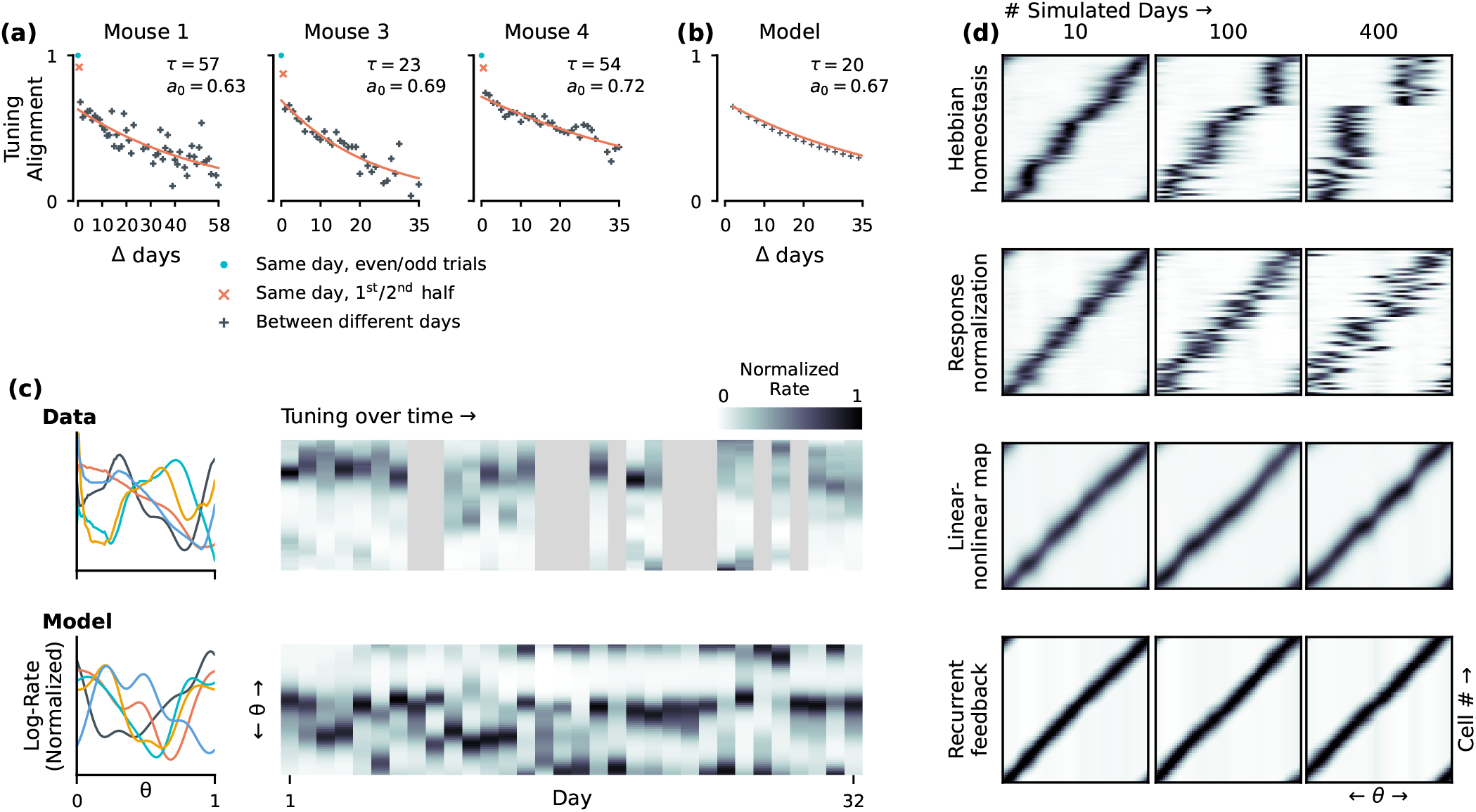
The statistics of drift observed in vivo are compatible with long-term stability. **(a)** The rate of drift can be measured using the cosine of the angle between two tuning curves (“alignment”), estimated on different days for the same cell (1: identical; 0: unrelated). Plots show tuning-curve alignment for a population of cells tracked for over a month, taken from three subjects in Driscoll et al. (47). Each point reflects the average alignment across the population for a pair of recordings separated by Δ days. Measurement noise exaggerates apparent tuning differences, so alignment was normalized via a bootstrap estimator such that tuning curves estimated from different trials on the same day were fully aligned (teal ‘o’). Alignment decays exponentially, with a similar timescales across subjects (“*τ*”). Extrapolating the day-to-day alignment (black ‘+’) to a separation of zero days (*a*_0_) does not yield perfect alignment, indicating that not all day-to-day tuning variability is explained by drift. This excess variability also cannot be attributed to systematic drift during the recording session (measured as the alignment between the first and second half of the recording session; red ‘×’). **(b)** We modeled tuning curves as samples from a log-Gaussian process over the latent space *θ*, with drift modeled as an Ornstein-Uhlenbeck random walk in tuning over time (Methods: *Simulated Drift*). We used a time-constant of *τ* = 45 days and applied *r* = 30% excess per-day variability to match the model to experimental data. **(c)** Tuning curves sampled from the model (bottom left) qualitatively resemble those in the experimental data. Tuning curves *in vivo*, however, exhibited nonuniform statistics in θ (top left). The statistics of drift (right) are also similar, with both model and data exhibiting day-to-day variability superimposed over slower long-term drift which exhibits punctuated stability. **(d)** Evolution of readout population tuning curves under simulated drift. The drift timescale and day-to-day variability were calibrated as in (b). All other parameters were the same as in Figure 4. Readouts with a stable internal model (“linear-nonlinear map” and “recurrent feedback”) exhibit long-term stability.

**Figure 8:**
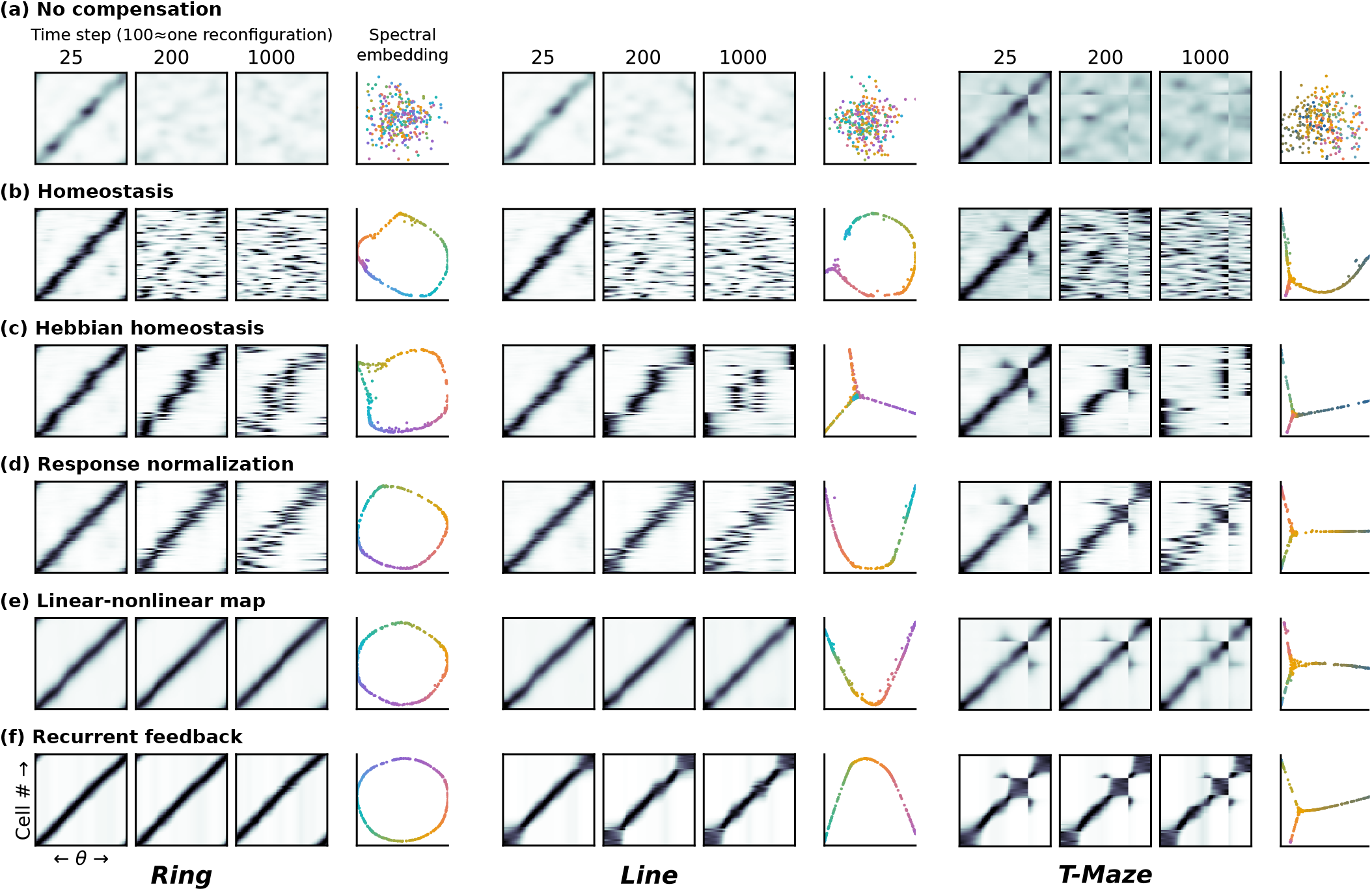
Self-healingplasticity stabilizes variousgeometries. We simulated representational drift under various self-healing plasticity scenarios as in Figure 4 of the main text, applied to different geometries. This figure illustrates a ring-shaped **(left)**, linear **(middle)**, and T-shaped **(right)** geometries for the encoded latent variable *θ*. The Gaussian-process synaptic activations for **x**(*θ*) were adapted to each geometry by shaping the covariance structure to match each geometry, keeping the correlation as a function of distance the same as in Figure 4 in the main text and Methods: *Simulated drift*. We simulated 1000 iterations of drift with time-constant *τ* = 100. Results are similar across all three topologies. Black-and-white plots show the configuration of the readout population code at various time-points, similarly to Figure 4 (left) in the main text. Colored plots show the result of applying unsupervised dimensionality reduction to the final readout population tuning curves (Python sklearn SpectralEmbedding (80); c.f. (43)). We applied this embedding to points sampled from five random ‘walkthroughs’ of *θ* with additive Gaussian noise *σ_ξ_* = 1.2 × 10^-2^ to emphasize the loss of signal-to-noise ratio in the absence of compensation. **(a)** Without compensation, the amount of variability in **y**(*θ*) that is related to *θ* decays, lowering the signal-to-noise ratio. Both the original tunings, and the capacity to encode *θ*, is lost. **(b)** With homeostasis, the original readout tuning curves are lost. However, homeostasis stabilizes the information-coding capacity of the readout. This is evidenced by the fact that nonlinear dimensionality reduction can still recover the underlying topology of *θ*. In this scenario, the readout **y**(*θ*) behaves much like the drifting encoding population **x**(*θ*). **(c)** Hebbian homeostasis provides some stability, but causes the readout population code to collapse around a few salient preferred *θ*_0_. **(d)** Response normalization compensates for the destabilizing impact of Hebbian homeostasis. However, noise causes readout neurons to swap their preferred tunings. **(e, f)** Long-term stability is possible in readouts with a stable internal model. Sharing of information among the readout population, modeled here as either a linear-nonlinear map or recurrent feedback, allows for more robust error correction.

**Figure 9:**
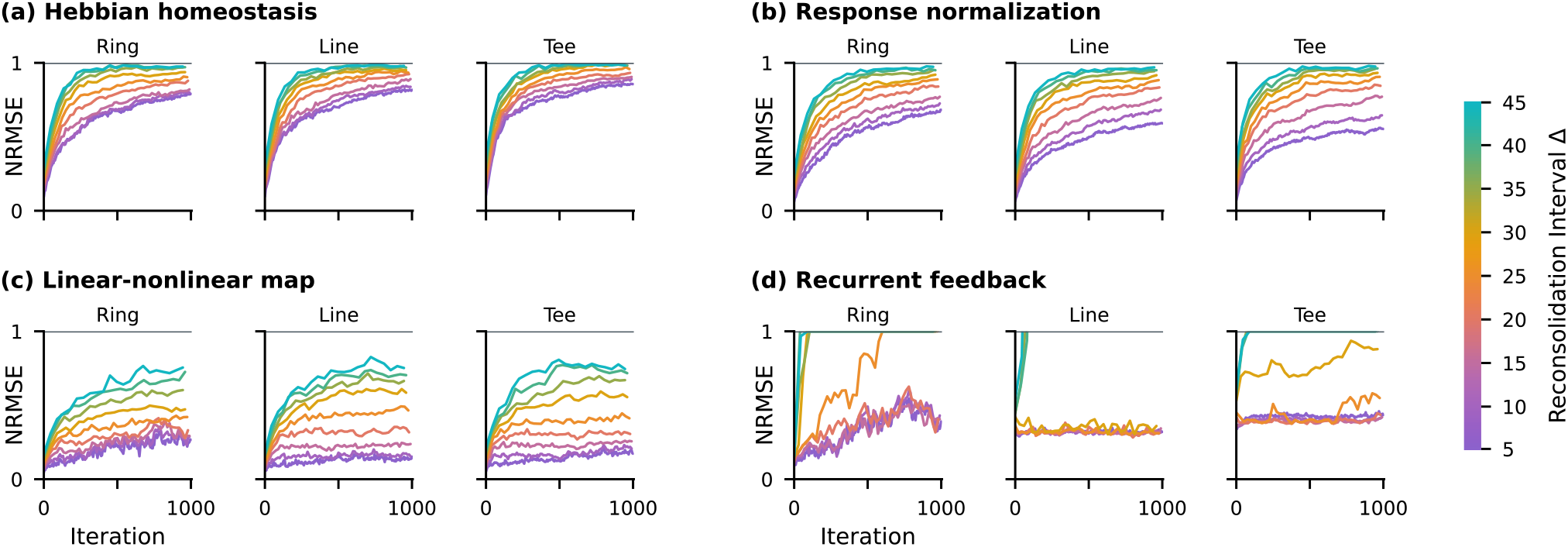
Larger amounts of drift between reconsolidation sessions reduces stability. Changing the rate of drift relative to the frequency of reconsolidation affects the stability of the readout population code. In these simulations, all parameters are the same as in Fig. 4 in the main text, with the exception of the frequency of reconsolidation Δ. In the main text, Δ = 5. We explore up to Δ = 45, equivalent to nine times faster drift. The rate of degradation for Hebbian homeostasis scales with the rate of drift, with **(a)** and without **(b)** response normalization. Error correction via linear-nonlinear map **(c)** behaves similarly. However, there is evidence for a stable steady-state solution for moderate rates of drift. The error of this steady-state solution increases with the drift rate. With recurrent feedback **(d)**, the population readout is stable for modest rates of drift, but loses stability above a certain rate (Δ ≈ 25).

## Code availability

Source code for all simulations is available online at github.com/michaelerule/selfhealingcodes.

## Acknowledgements

This work was supported by the European Research Council (Starting Grant F?EXNEURO 716643), the Human Frontier Science Program (RGY0069), the Leverhulme Trust (ECF-2020-352), and the Isaac Newton Trust.

## Supplemental Information

### Gaussian-process tuning curves

Equation (10) in the main text (Methods: *Simulated drift*) defines an Ornstein-Uhlenbeck (OU) random walk on the encoding weights **U**. This is equivalent to assuming that the synaptic activations of the encoding population undergo a random walk, and are samples from a stationary distribution of functions on *θ*.

Assume that individual encoding weights *u* vary randomly over time, sampling from a stationary distribution that can be approximated as Gaussian. Without loss of generality, choose units such that this distribution is a standard normal distribution 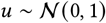. Denote the time-varying vector of synaptic weights for a single encoding neuron as **u**_*d*_. Now, consider the synaptic activation for an encoding neuron driven by features 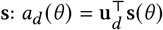. If 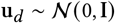, then the second moment ∑_*a*_(*θ*, *θ*′) = 〈*a_d_*(*θ*) *a_d_*(*θ*′)〉_*d*_ is:

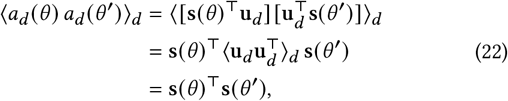

which is constant since **s**(*θ*′) do not change over time. A similar logic holds for the other moments, confirming that the OU drift on the encoding weights samples from a stationary distribution of activation functions.

If the encoding features **s**(*θ*) are sampled from a Gaussian process on *θ*, then OU drift on the encoding weights amounts to OU drift over a Gaussian-process distribution of activation functions. Let the encoding weights change as 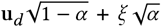, where *α* sets the drift rate and *ξ* are are the Gaussian noise sources as in Eq. 10 in the main text. Define the drift in synaptic activation at each time-point as 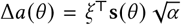. The updated synaptic activations *a*_*d*+1_(*θ*) are then:

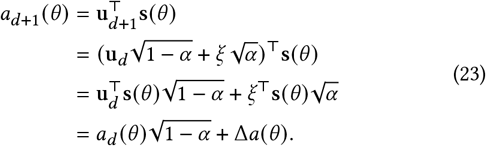

The increments Δ*a*(*θ*) are samples from a Gaussian Process 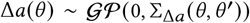, with second moment:

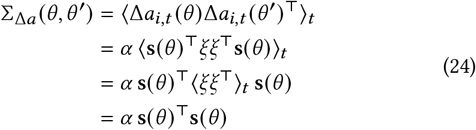

If units are chosen such that the encoding features **s**(*θ*) are zero-mean in *θ*, then the second moments in Eqs. (22–24) can be interpreted as covariances. The randomly-drifting encoding weights **u** are therefore equivalent to an OU random walk through the space of possible activation functions on *θ*. We use this to simplify computations.

If the synaptic activations *a*(*θ*) are samples from a stationary distribution of functions over *θ*, then the tuning curves *x*(*θ*) = *ϕ* [*a*(*θ*)] are also samples from a stationary distribution over possible tuning curves. Adding homeostasis scales and shifts the activation to achieve the target firing rate statistics, removing two directions of variability from this distribution. This regulates the amount of information about *θ* encoded in the firing-rate variations of the population **x**(*θ*) (Supplement: *Stability of encoded information*).

### Stability of encoded information

The model of drift described in Methods: *Simulated drift* and Supplement: *Gaussian-process tuning curves* conserves the information-coding capacity of **x**(*θ*). Because individual encoding neurons evolve independently in this model, the population of *N* encoding cells represents *N* independent samples from the distribution of tuning curves on *θ*. If we consider the large (*N*→*∞*) population limit, the average population variability related to *θ* is given by the expected variability caused by *θ* for the typical tuning curve:

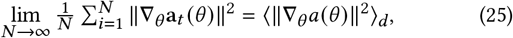

where 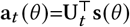 is the vector of synaptic activations for all encoding units at location *θ*. The expected amount of population variability driven by *θ* is conserved, and is a function of the covariance of the activation functions ∑_*a*_(*θ*, *θ*′):

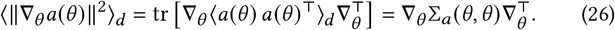

The second step in Eq. (26) follows from the linearity of the trace and the identity ||**q**||^2^ = tr[**qq**^⊤^]. The operator 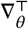 refers to differentiating ∑_*a*_(·, ·) in its second argument. This is in turn relates to the amount of variability in the features **s**(*θ*) that is driven by *θ*:

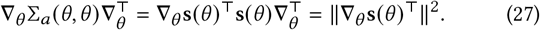

This shows that **a**(*θ*) inherits the correlation structure of **s**(*θ*), and that in the large population limit the variation in **a**(*θ*) driven by *θ* is approximately conserved.

Additional assumptions about the nonlinearity *ϕ*[·] are needed to show that stable information in **a**(*θ*) implies stable information in **x**(*θ*) = *ϕ* [**a**(*θ*)]. In the special case of an exponential nonlinearity *ϕ* = exp, the trace of Fisher information 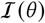 of 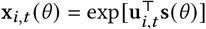 is proportional to the average variation in **a**(*θ*) driven by *θ*:

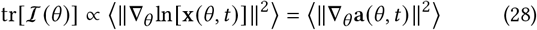

(Formally, the Fisher information is infinite when the noise in **x** is zero, but Eq. (28) can be viewed as the zero-variance limit of homogeneous and IID Gaussian noise with suitable normalization.) With a threshold nonlinearity, the dynamic range of each *a*(*θ*) must remain in a certain range to ensure that information is not lost due to the saturation in the firing-rate response. This can be ensured by homeostasis (24, 48, 79).

### Hebbian homeostasis as an emergent property

Here we explore a simplified linear model to make concrete the intuition that Eq. (5) in the main text should emerge through interactions between homeostasis and Hebbian learning. Consider a single (scalar) linear readout with inputs **x**, weights **w**, and output firing rate *y*:

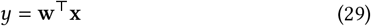

Let **X** and **Y** be a training dataset of presynaptic inputs **x** ∈ **X** and postsynaptic outputs *y* ∈ **Y**. Assume that *y* and **x** are both zero-mean over this dataset. Consider an Oja-style Hebbian learning rule of the form:

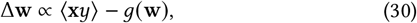

where *g*(**w**) represents stabilizing terms in the learning rule. Assume that learning has converged to steady state for some 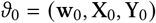, such that Δ**w** = 0. This implies that

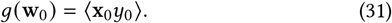

Now, consider incremental drift in the input encoding **X**_1_ = **X**_0_ + Δ_X_. This changes the readout’s firing to *y*_1_ = *y*_0_ + Δ_*y*_, where 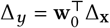 is small. This alters the activity statistics of *y*_1_, changing the firing-rate variance 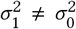. Assume that the variance of *y*_1_ has been restored by homeostatic processes that multiply the firing rate by a factor *γ* = *σ*_0_/*σ*_1_ = 1 + *δ*, where *δ* is a small parameter:

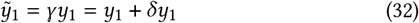

What is the influence of this new activity *ỹ*_1_ on plasticity? Evaluating Eq. (30) for (**x**_1_, *ỹ*_1_), and substituting in Eq. (31) yields:

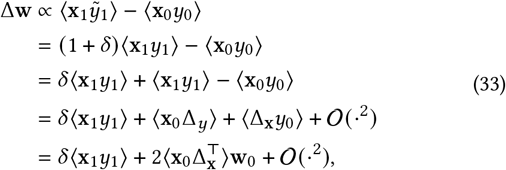

where 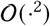 denotes all terms of second-order and higher in Δ_x_. If drift is uncorrelated with the current state, then 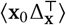 is zero to first-order in Δ_x_, and Eq. (33) simplifies to:

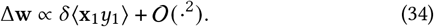

This is similar to the Hebbian term in Eq. (5) in the main text, if *δ ∝ ε_σ_* at first order. This is easily verified:

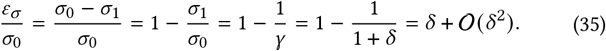

Eq. (35) is therefore equivalent to the Hebbian contribution to the Hebbian homeostatic rule in Eq. (5) in the main text, with *δ* = *ε_σ_*/*σ*_0_. What about the weight decay terms?

We assume that the norm of the weight vector, ||**w**||^2^, is conserved. Hebbian learning will generally disrupt this. If we assume that the norm of the weight vector is restored by weight decay −*ρ***w**, what value of *ρ* would keep the norm of the weight vector constant? Consider a weight update as in Eq. (35), with an unknown weight decay term −*ρ***w**:

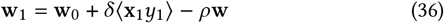

Assume that 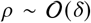. What value would *ρ* need to take to ensure that ||**w**_1_||^2^ = ||**w**_0_||^2^? The norm of the updated weight vector is:

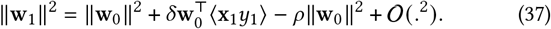

We see that ||**w**_1_||^2^ = ||**w**_0_||^2^ if 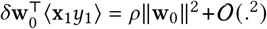, implying that

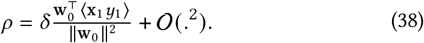

Since 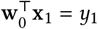, the term 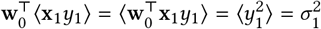 is equal to the firing-rate variability after perturbation by drift. This implies that

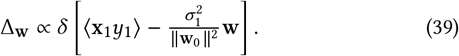

This suggests that the optimal value of weight decay is 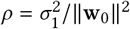. In practice we found that and setting *ρ* = 1 still led to good stability in simulations.

This derivation is not indented to prove that Hebbian homeostasis should arise in any specific physiological model, but rather to illustrate that a learning rule of this form might reasonably be expected to exist based on known Hebbian and homeostatic plasticity mechanisms.

### Weight filtering

The action of the Hebbian homeostatic rule (Eq. 5 in the main text) can be interpreted as a form of filtering. Consider a linear readout trained initially on day *d* = 0 with weights **W**_0_ (make dependence on *θ* implicit to simplify notation):

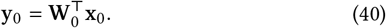

The encoding **x**_*d*_ changes on each day *d*. Tracking these changes entails translating the population code-words **x**_*d*_ into the code originally used on day 0. Using this translation 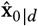, one might achieve an approximately stable readout.

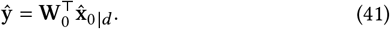

How could one estimate 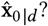 Consider estimating the code-words on day *d* from those on day *d* + 1. Let drift *d***x**_*d*_ be sampled from a known distribution 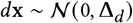. An estimate of 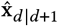 can be obtained via linear least-squares:

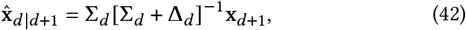

where **Σ**_*d*_ is the covariance of the code **x**_*d*_ on the previous day.

Eq. (42) provides the minimum squared error estimate of 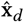. In the linear, Gaussian case this is also the Bayesian *maximum a posteriori* estimate. Applying Eq. (42) iteratively yields an estimate of the original code 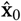, thereby translating the current representation **x**_*d*_ back through time to when the readout was first learned:

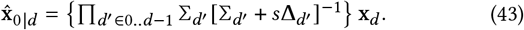

Now, consider the effect of Eq. (42) on a decoded **ŷ** by substituting Eq. (42) into Eq. (41):

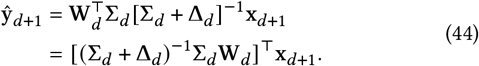

The expression [(Σ_*d*_ + Δ_*d*_)^-1^Σ_*d*_**W**_*d*_ in Eq. (44) is the same one used to re-train the readout from its own output (Eq. 15 in the main text, Methods: *Synaptic learning rules*, with **Δ**_*d*_ estimated as *ρ***I**):

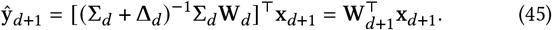

This illustrates that the Hebbian homeostatic learning rules outlined in the text can be viewed loosely as filtering the current code-words **x**_*d*_ to recover the original code **x**_0_ against which the readout was first trained. In a linear, Gaussian case the correspondence with Bayesian filtering is exact.

### Predictive coding as inference

In the main text (Results: *Predictive error feedback*), we argued that negative feedback of prediction errors removes variations in **y**(*θ*) that are inconsistent with a learned internal model. In particular, if recurrent weights **A**_*p*_ are learned through Hebbain learning, then this feedback selectively tracks only the modes of **y**(*θ*) known to enode information about *θ*. This leads the readout to infer a de-noised estimate 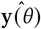 that can be used to update decoding weights. Here, we prove that this inference has an exact interpretation as Bayesian inference under certain circumstances.

Assume that the readout has internal states **z**, and has learned a prior for what values these states should take for encoding *θ*, Pr(**z**). This prior reflects the ground-truth distribution of **z** experienced when the inputs **x** are driven by true external inputs **s**(*θ*) during behavior. Assume that the error model for the feed-forwarded decoding **y**_*f*_ (Eq. 4 in the main text) in known, Pr(**y**_*f*_|**z**). For a given **y**_*y*_, the Baysian posterior estimate for 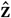 is

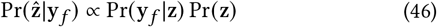

The estimate 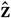 can be optimized by finding the posterior mode of Eq. (46). This can be done by maximizing the log-posterior:

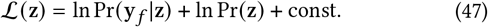

Now, consider the case where the prior on **z** is multivariate Gaussian. Let this prior be zero mean for convenience, without loss of generality, 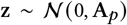. Let the observation model Pr(**y**_*y*_|**z**) be of the natural exponential family, where **z** are the natural parameters, and **y**_*f*_ are the natural statistics. Let *f*(·) be an element-wise function of **z** such that its derivative matches the firing-rate nonlinearity, *ϕ* (**z**) = *f*′(**z**). The log-prior and log-likelihood then take the forms:

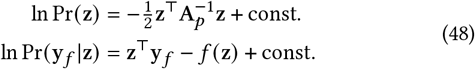

The states **z** can be optimized via gradient ascent of 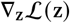 on Eq. (47), implying the following dynamics:

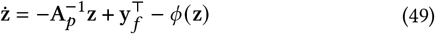

Multiplying through by the prior covariance **A**_*p*_ does not change the fixed points, and so Eq. (49) can be written as:

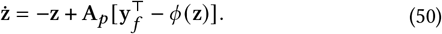

Eq. (50) is the same as Eq. (7) in the main text (Results: *Predictive error feedback*), with the exception of a time constant *τ_z_* which does not change the fixed points.

As an aside, modulating the feedback gain by a factor *κ* in Eq. (50) allows a neural population to dynamically adjust the influence of the external inputs and its internal model:

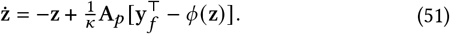

One might set *κ* to be small when one is confident in **y**_*f*_. This would cause learning to overwrite a learned **y**(*θ*) with external input. Conversely, larger *κ* can be used to rely on priors more heavily when **y**_*f*_ are uncertain. We conjecture that modulating the feedback gain might control whether internal models vs. external input dominate activity, and, as a result, synaptic plasticity.

Overall, the correspondence between predictive coding and Bayesian inference is exact when

1. Subthreshold activations **z** receive prediction-error feedback, and can be interpreted as the natural parameters of an exponentialfamily distribution for **ŷ** = *ϕ*(**z**).
2. Feedback weights **A**_*p*_ are proportional to the covariance of a Gaussian prior on **z**.
3. The nonlinearity implies a natural exponential family that can reasonably capture error and uncertainty in **y**_*f*_.

The natural exponential family includes most common firingrate models of neural dynamics, including linear-Gaussian, linear-nonlinear-Poisson, and linear-nonlinear-Bernoulli.

It is unlikely that this correspondence is exact *in vivo*. Symmetric Hebbian learning of the recurrent weights would potentiate correlations in **y** = *ϕ*(**z**), not **z**. This would lead the recurrent weights to learn the covariance structure in the firing rates, **A**_*p*_ ∝ Σ_*y*_, rather than synaptic activations **A**_*p*_ ∝ Σ_*z*_. These are only equal in the case of a linear readout. A saturating nonlinearity that approximates *ϕ*^-1^(·) when tracking pre-post correlations could compensate for this, but there is no experimental evidence for this.

Regardless, Eq. (50) generically restricts the activity in **y** to the *subspace* associated with encoding *θ* (Results: *Recurrent feedback of prediction errors*.). In large population codes for low-dimensional activity, this subspace is low rank. Eq. (50) is therefore likely to provide useful error correction via subspace projection, even if the matching of the amplitudes of specific modes of **y** to the internal model is inexact. The recurrent feedback model should therefore be interpreted as a qualitative hypothesis for how corrective feedback might aid in tracking drift. The precise interpretation of the feedback weights **A**_*p*_ is left to further studies.

### Learning as inference

Eq. (25) in the main text describes how the forward weights evolve based on the correlation between a prediction error **ε** = **ŷ** – **y**_*f*_ and the inputs **x** (Methods: *Population simulations*). It also includes a weight decay term −ρ**W**. This trains the forward weights to minimize prediction errors, subject to the constraints that the weights should not grow too large. Analogously to how negative feedback optimizes a trade-off between input and an internal estimate of **ŷ**, this optimization corresponds to Bayesian inference when certain quantities are equated to the parameters of a distribution from the natural exponential family.

Consider training the readout weight vector **w** for a single decoding unit, using *L* training examples, each consisting of an input **x** and a desired output *y*. The readout makes predictions *y_f_* = *ϕ*(*z*), where *z* = **w**^⊤^**x**. Now, define a zero-mean Gaussian prior for these weights, 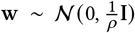. Let the firing-rate nonlinearity *ϕ*(*z*) = *f*′(*z*) correspond to the derivative of a known function *f*(·), and assume that our observation model is taken from a natural exponential family with natural parameter *z* and natural statistics *y*. Up to constants, the log-posterior for **w** can then be written as

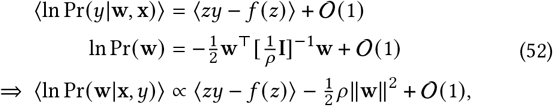

where the expectation 〈·〉 averages over the training data. This objective can be optimized by ascending the gradient **▿_w_** of the loglikelihood, implying to the following weight dynamics:

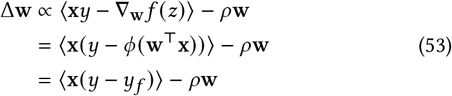

Eq. (53) is equivalent to the error-based learning rule used in the main text (Eq. 25), if one applies this update in discrete steps with learning rate *η* to a population readout.

As was the case for prediction-error feedback, this statistical interpretation is unlikely to correspond exactly to processes *in vivo*. Nevertheless, learning rules resembling Eq. (53) do emerge in some timingdependent plasticity rules (78). In this work, such rules are used primarily to initialize the readout weights **W** or recurrent weights **A**_*r*_ for the linear-nonlinear map. Subsequent evolution of **W** is then assumed to follow the unsupervised Hebbian homeostatic rule in Eq. (5) in the main text. A biologically plausible interpretation of supervised, error-driven learning is therefore not essential to the core results regarding the tracking of representational drift.

### Calibrating the model to data

The results in the main text are abstract, because drift statistics vary across brain areas, species, and experimental conditions. Stability depends on many unknown parameters, include drift rate, noise levels, population-code redundancy, and the frequency of reconsolidation. Nevertheless, in Supplemental Figure 4, we present a best-effort calibration of our simulations to the Driscoll et al. (2017) data recorded from mouse posterior parietal cortex (2, 47).

We calibrated a model of simulated drift using three subjects from Driscoll et al. (47) (mice M1, M3, and M4). For each subject, we selected a subset of fifteen recording sessions sharing common neurons (N=10, 60, and 83 neurons from M1, M3, and M4 respectively). Each subpopulation was tracked for over a month (58, 35, and 38 days respectively), with modest gaps between consecutive recording sessions (no more than 13, 8, and 10 consecutive missing days, respectively). For each day, we filtered log-Calcium fluorescence traces between 0.03 and 0.3 Hz and normalized them by z-scoring. We aligned filtered traces from successful trials to task pseudotime (“*θ*”) based on progress through the maze. These traces were averaged to estimate neuronal tuning curves.

The tuning curves can be interpreted as vectors in a highdimensional space, with a different component for each location *θ*. We used the cosine of the angle *ψ* between two tuning curves as an “alignment” measure to quantify drift. This can be computed as the average product 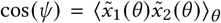 between normalized (z-scored) tuning curves 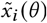, and is 1 if the tuning curves are identical and 0 (on average) if they are unrelated.

This estimator is biased by measurement noise and trial-to-trial variability. We compensated for this by bootstrap-resampling the average alignment between two tuning curves from the same cell and day, where each tuning curve is estimated as an average over a different random subsets of trials. This baseline was calculated separately for each neuron and day, averaged over ten random samples. Bias was removed by dividing the tuning-curve alignment measure by this baseline. This retains excess per-day variability that cannot be explained by drift, noise, or trial-to-trial fluctuations.

Alignment decayed exponentially over time (Figure S4-a). The decay time constant (*τ*) indicates the drift rate. The extrapolated value at Δ = 0 days (*a*_0_) indicates the excess per-day variability. We estimated these parameters using least-squares exponential regression. Results were similar across subjects (*τ* = 57, 23, 54 and *a*_0_ = 0.63, 0.69, 0.72; for M1, M3, M4 respectively). We used the average values of these parameters across subjects (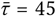 and 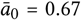) to calibrate a model of drift (Figure S4-b). This calibration does not provide all the information needed for a principled comparison between simulations and experiments. However, it seems reasonable to conjecture that among the ~10, 000**+** synaptic inputs to a given pyramidal cell, there is sufficient redundancy to recover a readout with accuracy comparable to the 100 idealized encoding units considered here.

Comparing simulation and experiment also requires assumptions about the frequency of “self-healing” reconsolidation, relative to the drift rate. In our simulations we apply self-healing every Δ = 5 time points. If we assume that “self-healing” occurs once per day *in vivo*, then an excess per-timepoint variability of *r* = 30% and a time constant of *τ* = 45 days matches the simulated drift to the Driscoll et al. (2017) (2) data. This implies approximate stability out to ~10 days using redundancy alone (fixed readout weights), a few months using Hebbian homeostasis, and over a year if the readout contains a stable internal model (Figure S4-d). However, a rigorous test of how these ideas apply *in vivo* would require new experimental studies.

### Normalized Root Mean-Squared Error (NRMSE)

In Supplemental Figures 1, 2, 3, and 6 we summarized the stability of the readout population code by measuring the normalized distance between the initial, trained readout firing-rates **y**(*θ*), and the firing rates on a given time-step **y**_*d*_(*θ*).

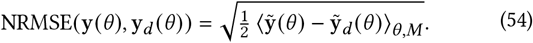

The values **ỹ** and **ỹ**_*d*_ reflect normalized tuning curves, in which the firing-rate function *y*(*θ*) for each readout neuron has been z-scored The average 〈·〉_*θ,m*_ is taken over all *M* decoding units and all *L* values of *θ*. The normalization by 1/2 ensures NRMSE ranges from 0 (identical codes) to 1 (chance level).

## Notes

### Competing Interest Statement

The authors have declared no competing interest.

### Summary of Updates

We have updated the manuscript to adjust emphasis and improve clarity.

